# Structural basis of substrate recognition for proteasome degradation by prokaryotic ubiquitin-like protein ligase PafA

**DOI:** 10.64898/2026.04.10.717840

**Authors:** Alicia Plourde, Adwaith B. Uday, Taylor J. B. Forrester, Natalie Zeytuni, Siavash Vahidi

**Author notes:** **Correspondence to:** Siavash Vahidi; or Natalie Zeytuni. These authors contributed equally to this work (Alicia Plourde, Adwaith B. Uday).

## Abstract

Selective protein degradation in some bacteria is performed by the Pup-proteasome system, in which the ligase PafA tags hundreds of substrates for proteasomal degradation. How a single enzyme achieves such broad substrate specificity in the absence of conserved sequence motifs has remained unclear. Here we determine structures of PafA in complex with a pupylated substrate and show that substrate recognition is mediated by a minimal and highly distributed interface. PafA samples an ensemble of closely related conformations that collectively position the target lysine residue for modification. This recognition mechanism arises from a combination of structured contacts and dynamic elements on both the enzyme and substrate, enabling geometric compatibility rather than sequence-specific interactions. These findings reveal an ensemble-driven mechanism of molecular recognition that explains how broad substrate specificity is achieved and provides a framework for understanding selective protein degradation in prokaryotes.

## Introduction

Regulated protein degradation is essential across all domains of life. In eukaryotes, selective protein turnover is primarily performed by the ubiquitin-proteasome system (UPS), in which ubiquitin is covalently attached to lysine residues on target proteins, thereby marking them for proteasomal degradation (*1–4*). Bacteria within the Nitrospirota and Actinomycetota phyla encode a functionally analogous Pup-proteasome system (PPS) (*5–7*). Importantly, the PPS is essential for maintaining proteome integrity and supports the survival of *Mycobacterium tuberculosis* during infection (*6, 8, 9*). In the PPS, Pup (<u>p</u>rokaryotic <u>u</u>biquitin-like <u>p</u>rotein) functions analogously to ubiquitin and is ligated post-translationally to target proteins by PafA (<u>p</u>roteasome <u>a</u>ccessory <u>f</u>actor <u>A</u>) via a process termed pupylation (fig. S1) (*7, 10*). In several species, Pup is expressed with a C-terminal glutamine (Pup^Q^) and must be deamidated by Dop (<u>D</u>eamidase <u>o</u>f <u>P</u>up) to produce a ligation-competent Pup^E^ (*11–13*). PafA ligates the C-terminal glutamate side chain of Pup^E^ to an exposed lysine side chain on a target protein via a two-step ATP-dependent mechanism (*14*), forming an isopeptide bond (fig. S1). Once pupylated, the target protein can be either recognized by the proteasome regulatory particle (RP) and degraded, or it can be depupylated by Dop and therefore rescued (*15*).

Although the PPS and the UPS both rely on covalent substrate tagging, they employ fundamentally distinct enzymatic mechanisms mediated by structurally unrelated proteins. In the UPS, substrate specificity is distributed across hundreds of E3 ubiquitin ligases, each responsible for modifying a narrow subset of targets (*1–4*). In contrast, PafA is the sole Pup ligase which consolidates the roles of the E1 ubiquitin-activating enzyme, E2 ubiquitin-conjugating enzyme, and E3 ubiquitin ligase into a single enzyme, and is responsible for tagging hundreds of pupylation targets (*4, 5, 7, 10, 16*). Strikingly, these pupylation targets share no discernible sequence motifs or structural features, preventing reliable prediction of modification sites (*17, 18*). The broad substrate tolerance of PafA has also been harnessed as a proximity-labelling tool to probe protein-protein interactions in eukaryotic systems, underscoring its ability to engage non-native proteins with little apparent specificity (*19*). Despite pupylating a diverse set of targets for proteasome degradation, the structural basis by which PafA recognizes and engages its protein substrates remains incompletely defined.

Although X-ray crystal structures of PafA have been determined in isolation and in complex with Pup (*16, 20*), they do not reveal how PafA discriminates among target proteins. Meticulous site-directed mutagenesis combined with biochemical assays by several groups have implicated several surface features in substrate recruitment, including an extended, positively charged loop adjacent to the active site and additional basic surface patches that are proposed to interact with acidic residues on target proteins (*21, 22*). Despite extensive biochemical characterization, the molecular mechanism underlaying the broad substrate specificity of this single enzyme in the absence of conserved sequence motifs remains elusive. Here, we determine structures of PafA in complex with a pupylated substrate and show that substrate selection is mediated by a sparse and distributed interface that encodes specificity through three-dimensional surface geometry rather than primary sequence. Together with published biochemical data (*20–22*) and those introduced here, our work helps define the molecular basis of substrate engagement by PafA and provides a framework for understanding how broad substrate specificity can be achieved in the absence of conserved recognition motifs.

## Results

### Domain swapping occludes the active site and is relieved by engineering a monomeric PafA

Two X-ray crystal structures of PafA have been reported to date. In one structure, the biological assembly contains a domain-swapped dimer [PDB: 4B0T (*20*)], whereas the other captures a monomeric PafA bound to Pup [PDB: 4BJR (*16*)]. In the domain-swapped PafA_dimer_ structure, the N-terminal β1/α1 motifs are exchanged between subunits through a short ^50^SS^51^ hinge immediately following α1 (fig. S2, A and B). This domain swap generates extensive secondary inter-subunit contacts absent in the monomeric structure, burying ∼6300 Å^2^ of surface area (*23*), with the C-terminal domain (CTD) of one subunit packing against the active-site β-sheet cradle and extended loop of the opposing subunit (fig. S2, D and E) (*20, 21*). Notably, the CTD residue R447 interacts with E18 of β1 and with E70’ of β4, both from the opposing intertwined active site (fig. S2D). Moreover, PafA and Dop are close structural homologues (RMSD 1.9 Å, Z-score 15.3), yet Dop is consistently observed as a monomer in all available structures (fig. S2C) (*13, 20, 24, 25*). Interestingly, the short ^50^SS^51^ hinge that mediates domain swapping in PafA aligns with the position of a ∼40-residue loop unique to Dop, suggesting that increased flexibility in this region disfavours domain swapping and stabilizes the monomeric state.

We recently showed that the domain-swapped PafA_dimer_ forms in solution, with an interface consistent with that observed in the crystal structure (*26*). The subunit arrangement in PafA_dimer_ occludes the active site and outcompetes low-affinity target proteins (*26*), suggesting that a monomeric PafA construct would favour productive target protein engagement. Accordingly, we aimed to engineer variants that promote monomer formation by targeting the ^50^SS^51^ hinge and the inter-subunit secondary contacts within PafA_dimer_. Size-exclusion chromatography (SEC) showed that inserting a three-residue linker in the hinge (PafA_hinge_: ^51^S-<u>SGG</u>-N^52^) or disrupting the secondary contacts involving R447 that stabilize PafA_dimer_ using a PafA_R447G_ variant, effectively monomerized PafA (Fig. 1A). We next compared the pupylation activity of PafA_hinge_ and PafA_R447G_ with wildtype PafA_monomer_ (isolated by SEC) using PanB, a well-characterized and physiologically relevant pupylation substrate that catalyzes an early committed step in pantothenate (vitamin B_5_) biosynthesis in mycobacteria, as the target protein (*14, 24, 27*). We reconstituted the pupylation system *in vitro* and monitored the formation of the PanB∼Pup^E^ product over time by SDS-PAGE followed by band densitometry. Relative to PafA_monomer_, PafA_hinge_ was ∼40% less active, whereas PafA_R447G_ was moderately more active than the wildtype enzyme (Fig. 1, B and C). These results establish a catalytically competent monomeric PafA_R447G_ variant that is suitable for structural characterization of the PafA-substrate complex.

**Figure 1.**
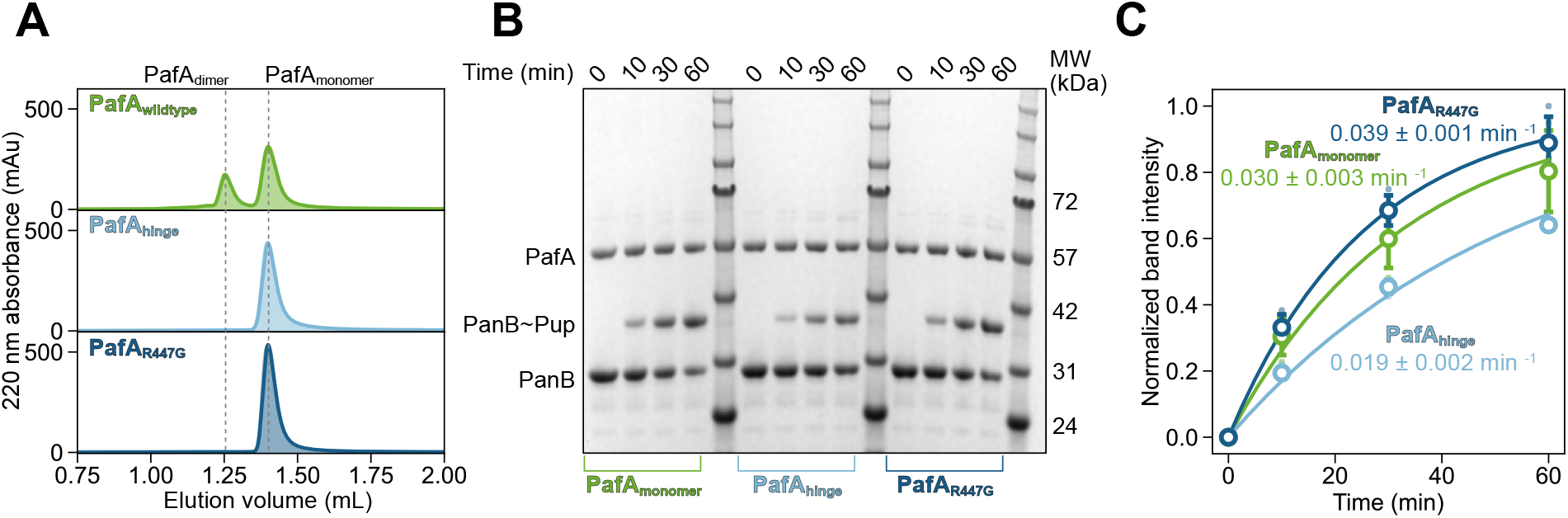
Monomerization of PafA yields an active enzyme. **A.** HPLC-based SEC traces of 10 µM PafAwildtype, PafAhinge, and PafAR447G. PafAwildtype (green) has two distinct peaks, corresponding to the domain-swapped dimer and monomeric fractions. In contrast, both PafAhinge (light blue), and PafAR447G (dark blue) variants elute as a uniform monomeric peak. **B**. The pupylation system was reconstituted *in vitro* with an isolated sample of wildtype PafAmonomer (left side – green), PafAhinge (middle – light blue), and PafAR447G (right – dark blue), and monitored via SDS-PAGE. As the reaction proceeds, the PanB band depletes in intensity and the formation of pupylated PanB (PanB∼Pup) increases. **C**. SDS-PAGE-based densitometry of the intensity of the PanB∼Pup band relative to the PafA band at the reaction onset. The individual data points (filled circles) and the mean values (open circles) from three technical replicates are shown, with error bars indicating the standard deviation across replicates. Wildtype PafAmonomer pupylates PanB at a rate of 0.030 ± 0.003 min ^-1^, whereas PafAhinge catalyzes the reaction at 0.019 ± 0.002 min ^-1^, and PafAR447G pupylates PanB the fastest at 0.039 ± 0.001 min ^-1^.

### Isolation of the PafA-ATP-PanB∼Pup^E^ quaternary complex

Formation of a stable PafA-ATP-PanB∼Pup^E^ quaternary complex is essential for determining its structure. Our strategy for complex preparation was to generate a partially pupylated PanB substrate and subsequently trap a product-bound PafA complex under turnover-inhibited conditions. PafA catalyzes pupylation via a two-step mechanism involving formation of a phosphorylated Pup intermediate at the active site, where ATP binds on one side of the catalytic cradle and the Pup C-terminal glutamate enters from the opposite side, positioning the substrate lysine for nucleophilic attack (figs. S1 and S2) (*14*). PanB assembles as a decamer by stacking two pentameric rings face-to-face, stabilized by the swapping of C-terminal α-helices between opposing subunits (*28, 29*). PafA binds Pup^E^ with ∼10-fold higher affinity than PanB (*K*_D_ ∼2 and ∼20 µM, respectively), and binds the PanB∼Pup^E^ product with affinity comparable to that for free Pup^E^ (*14, 16, 20, 30, 31*). We reasoned that converting PanB into the PanB∼Pup^E^ conjugate would shift binding toward a tighter, longer-lived PafA-bound state, thereby improving our ability to trap a homogeneous quaternary complex for high-resolution structural studies. Accordingly, we reconstituted the pupylation system *in vitro* with 1 µM PafA_R447G_ and 80 µM of each PanB (subunit concentration) and Pup^E^ in a 10-mL reaction volume to drive the formation of PanB∼Pup^E^ to high yield. We purified the PanB∼Pup^E^ product from lower molecular weight species by SEC. SDS-PAGE band densitometry indicated that approximately 40% of PanB subunits were pupylated (fig. S3). Assuming random pupylation across the 10 subunits of the PanB decamer (*p* = 0.40 per subunit), the fraction of PanB decamers carrying at least one Pup^E^ is 1 - (1 - *p*)^10^ = 1 - 0.6^10^ ≈ 99%. We incubated 200 µM PanB∼Pup^E^ (PanB subunit concentration) (fig. S3) with 0.5 mM ATP·Mg and 175 µM catalytically inactive PafA_D64N/R447G_ to prevent any further substrate turnover during complex assembly. We next used SEC to isolate the assembled product-bound quaternary complex for single-particle electron cryomicroscopy (cryo-EM) data collection (Fig. 2A).

**Figure 2.**
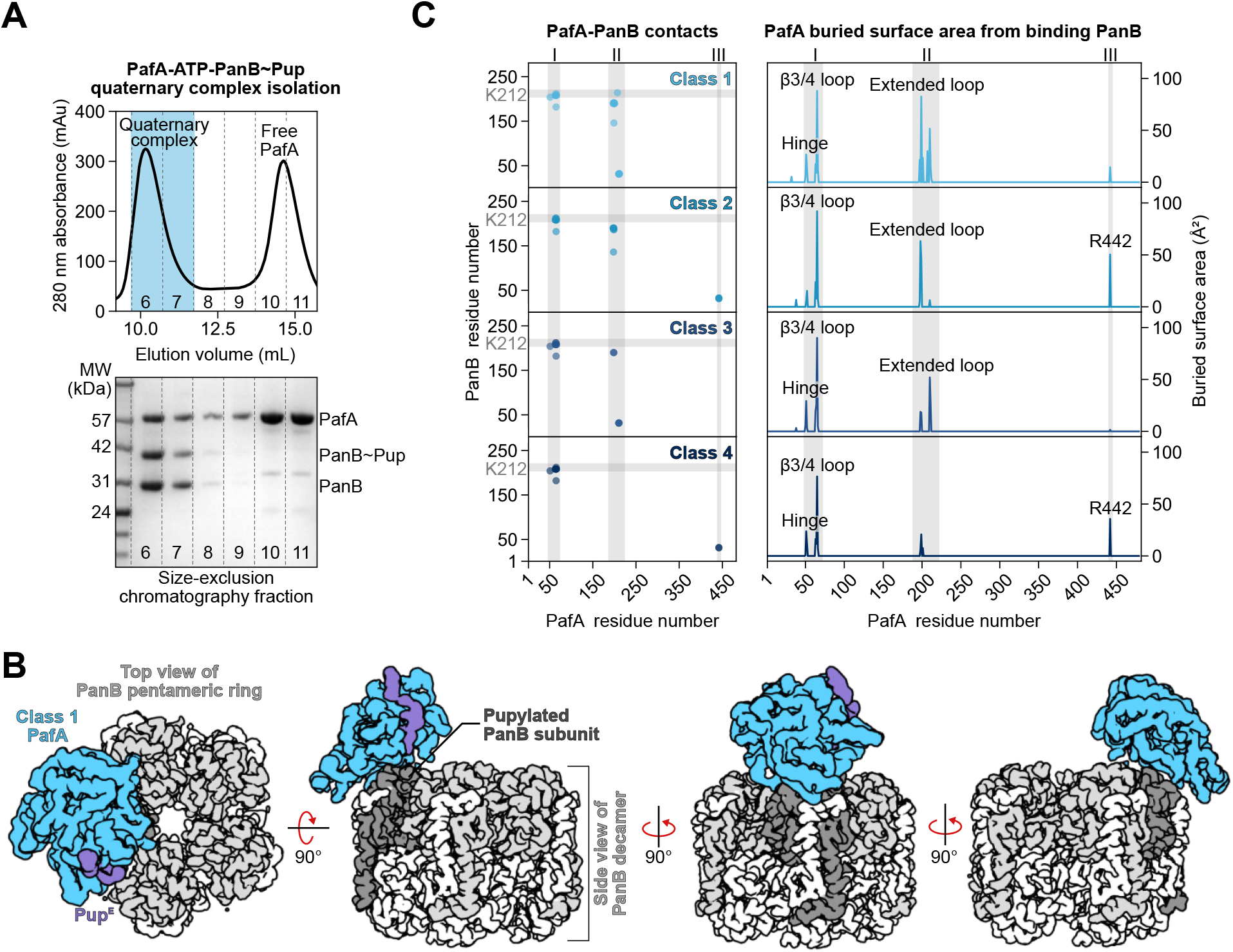
Architecture of the PafA-ADP-PanB∼Pup quaternary complex. **A.** The PafA-ATP-PanB∼Pup complex was assembled and purified via size-exclusion chromatography. SDS-PAGE analysis of the quaternary complex shows bands corresponding to the molecular weights of PafA (55 kDa), PanB∼Pup (36 kDa), and PanB (30 kDa). **B**. Cryo-EM map of the PafA-ADP-PanB∼Pup quaternary complex class 1. Pup (purple) occupies the Pup-binding groove of PafA (light blue), while PafA engages the apical surface of PanB (grey). **C**. Left – Residues of PafA (x axis) and PanB (y axis) within 5 Å are shown for each refined model. The region ±5 residues around the pupylated Lys212 in PanB is indicated by a grey horizontal bar. Three regions of PafA contacting PanB (I-III) are indicated by grey vertical bars. Region I corresponds to the ^50^SS^51^ hinge and the β3-4 loop (residues 63-66), region II is the extended loop (residues 196–216), and region III corresponds to R442 from the C-terminal domain. Right – Buried surface area per residue of PafA at the PafA-PanB interface is displayed for each model, with regions I-III highlighted in grey.

### A sparse, distributed interface underlies PafA substrate recognition

Our cryo-EM data processing pipeline (fig. S4) identified multiple particle populations, including assemblies with one, two, or more PafA molecules bound per PanB decamer (fig. S4). Particles corresponding to complexes with a single PafA bound were selected for downstream processing. These particles were refined using non-uniform refinement with C1 symmetry to generate an initial consensus reconstruction. Focused classifications on the PafA binding region further separated distinct conformational states, yielding four classes that were independently refined. Focused refinement was subsequently performed on these classes separately to improve the density at the PafA-PanB interface. Final maps were generated by combining the maps from the focused refinements, resulting in four reconstructions used for molecular modelling (Fig. 2B, figs. S5 and S6).

The resulting maps reached near-atomic resolution for the PanB core (2.5 – 4.0 Å), with reduced local resolution observed for PafA (4.0 – 6.5 Å) (table S1), suggesting that PafA is not tightly coupled to PanB and likely samples a continuum of closely related conformations, as discussed below. In the refined models, all components adopt their expected globular folds. PanB assembles into a decamer of two stacked pentameric rings, with each subunit forming a modified (β/α)_8_-barrel extended by a C-terminal α9 helix that mediates a domain swap to stabilize the decameric assembly (fig. S7). The primary pupylation site, K212 (*31*), is positioned on helix α8 at the apical surface of PanB. Structural alignment with the previously reported PanB X-ray structure [PDB: 1OY0 (*28*)] yields a Cα RMSD of ∼1 Å, indicating close agreement. PafA is monomeric, as expected, and adopts its characteristic fold, comprising a large N-terminal domain (residues 1-410) and a smaller CTD (residues 411-482), with a Cα RMSD of ∼1.7 Å compared to the PafA-Pup_Δ1-37_ fusion crystal structure [PDB: 4BJR (*16*)] (fig. S8). The PafA-Pup^E^ interface within the quaternary complex closely resembles that observed in the PafA-Pup_Δ1-37_ crystal structure [PDB: 4BJR (*16*)], burying an average of ∼1200 Å^2^ of solvent-accessible surface area across the four quaternary complex models (*23*). Although the quaternary complex was isolated in the presence of ATP, the electron density supported only ADP, with no resolved density for the γ-phosphate. Accordingly, ATP was not modeled in the final structure. This may reflect ATP hydrolysis by residual active enzyme during sample preparation, but it could also arise from flexibility or disorder of the γ-phosphate. The present data do not allow us to distinguish between these possibilities.

How PafA selects specific lysine residues for pupylation remains poorly understood and prior bioinformatic analyses have failed to identify any consensus sequence or structural motif (*17, 18, 22, 32*), suggesting that substrate recognition is governed by higher-order structural features rather than primary sequence determinants. Our structural models provide an explanation for this long-standing gap. PafA engages PanB through a minimal and highly distributed set of contacts, burying only 220-420 Å^2^ of solvent-accessible surface area (Fig. 3A). This interaction is organized around the Pup^E64^-PanB^K212^ isopeptide bond (Fig. 3A – inset) while multiple, spatially separated regions of PafA make contacts with the apical surface of PanB. The observed geometry of this interface is consistent with a catalytically competent substrate-bound state, as the target lysine is positioned adjacent to the location of the Pup C-terminal glutamate observed in one of the published structure of PafA [PDB: 4BJR (*16*)], supporting a configuration compatible with nucleophilic attack on the phosphorylated Pup intermediate. In the pre-reaction complex, formation of the phosphorylated Pup intermediate and proximity of the reactive lysine would further introduce favourable electrostatic interactions, thereby effectively increasing the buried surface area and stabilizing catalytically productive configurations. These regions include the ^50^SS^51^ hinge and the β3/4 loop (region I), the extended loop (region II), and CTD residue R442 (region III) (Fig. 2C, Fig. 3, figs. S9-11). Across the four refined quaternary models, the structural elements highlighted in these regions engage PanB to varying extents, producing distinct contact patterns and differences in buried surface area (Fig. 2C, table S2). This variability suggests a flexible binding mode rather than a rigidly defined interface (movie S1).

**Figure 3.**
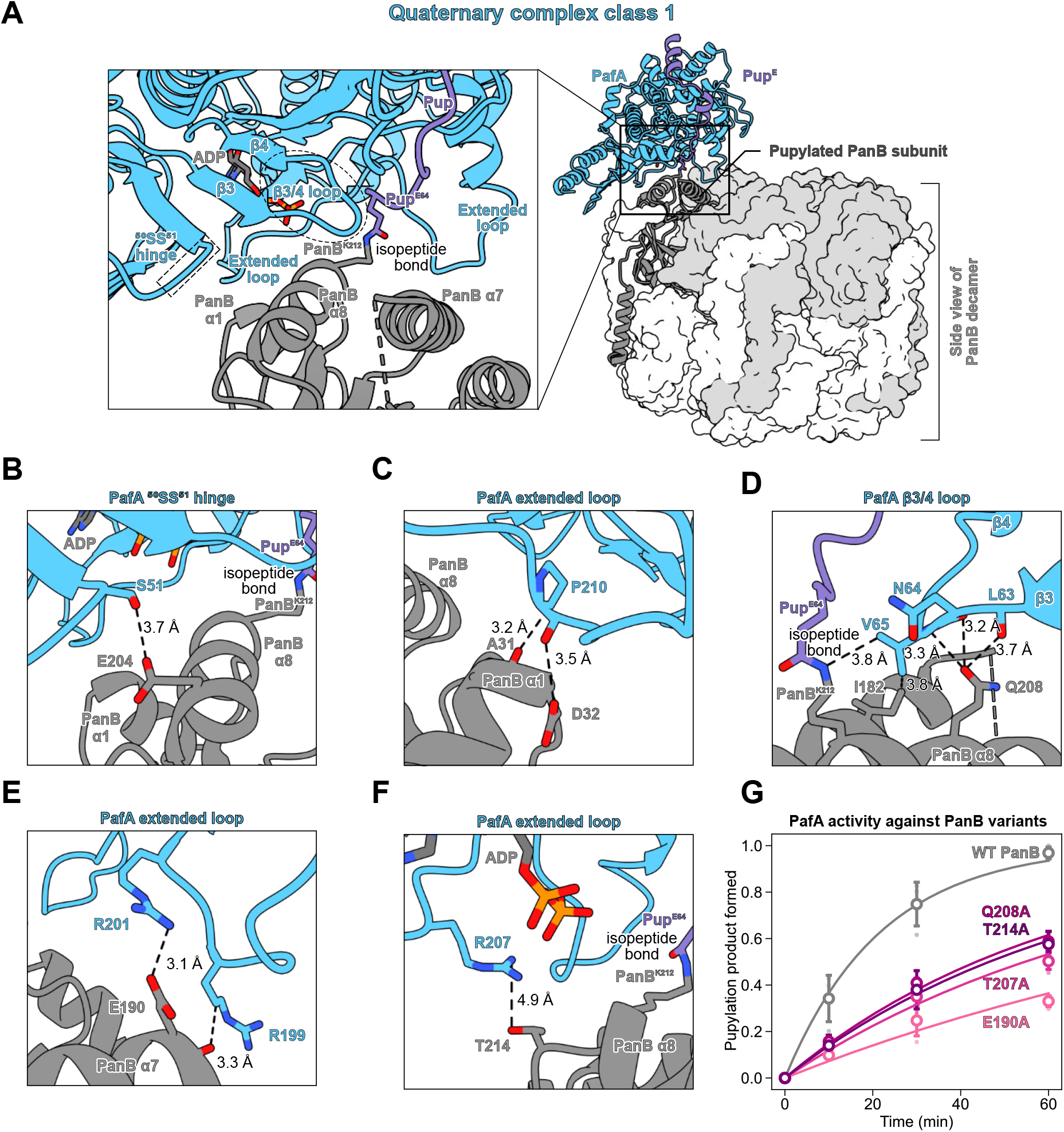
PafA engages the apical surface of PanB. **A.** The refined model of the class 1 quaternary complex is shown. PafA (blue), Pup (purple), and the pupylated PanB subunit (dark grey) are depicted as cartoons, while the remaining nine PanB subunits are rendered as a surface. PafA engages Pup within the Pup-binding groove and simultaneously contacts the apical surface of the pupylated PanB subunit. The inset highlights the PafA-PanB interface, with the three apical helices of PanB (α1, α7, and α8, the latter containing the pupylated K212) indicated. The isopeptide bond between PanB K212 and Pup E64 is shown, and PafA’s ^50^SS^51^ hinge, β3-4 loop, and extended loop involved in target protein engagement are highlighted. For clarity, PafA α1 (residues 33-46) is omitted from the inset. **B**. Interaction between PafA S51 and PanB E204, 3.7 Å (Oγ-Oε1). **C**. Interactions between PafA P210 and PanB A31, 3.2 Å (CB-O), and PafA P210 and PanB D32, 3.5 Å (O-Oδ1). **D**. Interactions between PafA L63 and PanB Q208, 3.7 Å (O-Oε1), PafA D64N and PanB Q208, 3.2 Å (O-Oε1), PafA V65 and PanB I182, 3.8 Å (Cγ1-Cδ1), PafA V65 and PanB Q208, 3.2 Å (N-Oε1), and PafA V65 and PanB K212, 3.8 Å (Cγ 2-NZ). **E**. Interactions between PafA R199 and PanB E190, 3.3 Å (Cγ-O), and PafA R201 and PanB E190, 3.1 Å (NH2-Oε2). **F**. Interactions between PafA R207 and PanB T214, 4.9 Å (NH1-Oγ1). **G**. PafA activity assay on PanB variants. The mean values (open circles) from three technical replicates are shown, with error bars indicating the standard deviation across replicates. PafAR447G was used to catalyze the pupylation reaction for wildtype PanB (grey) and four PanB variants to compare their pupylation rate relative to wildtype PanB: E190A (light pink – 0.17 ± 0.03), T207A (medium pink – 0.28 ± 0.04), Q208A (magenta – 0.35 ± 0.04), and T214A (purple – 0.33 ± 0.03).

Across all models, the majority of PafA-PanB contacts are formed with a single PanB subunit, with only limited contributions from neighbouring subunits. Region I provides a central set of contacts near the pupylated K212 on PanB. S51 from the PafA hinge interacts with PanB E204 (α8) in classes 1, 3, and 4, with the closest interaction observed in class 4, where S51 Oγ forms a hydrogen bond with E204 Oε1 at 3.1 Å (Fig. 3B and fig. S11C). The adjacent β3/4 loop of PafA contains residues previously implicated in catalysis and activity by the Darwin, Weber-Ban, and Gur groups: D64, which has been proposed to act as a catalytic base during pupylation (*20, 33*), and V65, where substitution to serine reduces pupylation efficiency (*21*). Consistent with their functional importance, both residues contact the α8 helix of PanB in our models. In our catalytically inactive construct (PafA_D64N_), N64 forms a hydrogen bond with PanB Q208 (3.0 Å in class 4; 3.2 Å in class 1), while V65 engages in close packing with PanB I182 (α7), Q208 (α8) and the pupylated K212 (α8) (Fig. 3D, figs. S9C, S10C, and S11D). In region II, PafA’s extended loop further reinforces the PafA-PanB interaction through a network of electrostatic contacts. In the first three classes, the extended loop residues of PafA T197, T198, R199, and R201 are positioned towards PanB E190 (α7), including a salt bridge between PafA R201 and PanB E190 at 3.1 Å in class 1 (Fig. 3, E and F, and figs. S9D, and S10D). In addition, R199 is positioned near the C terminus of PanB helix α6, where it engages the helix dipole and contributes a favourable electrostatic interaction. A disordered loop from a neighbouring PanB subunit (residues ∼160-175) is consistently observed close to the PafA surface. Although not explicitly modeled due to the lack of well-resolved density, this region may contribute additional transient contacts, suggesting that disordered elements could further stabilize the complex through dynamic interactions. Additional contacts include packing between PafA R207 and PanB T214, as well as interactions between PafA P210 and the α1 helix of PanB in two models (Fig. 3 C and F, and fig. S10B). Region III, corresponding to R442 from the CTD, positions the guanidinium group of R442 near the backbone carbonyl of PanB D32 on helix α1 in classes 2 and 4 (figs. S9A and S11B). R442 engages the helix dipole of PanB α1, contributing a favourable electrostatic interaction that is further reinforced by the distribution of charged residues along this helix, including D32 at the C terminus and R28, H24, H23, and R22 positioned along its length, which collectively amplify the local electrostatic field. Taken together, the PafA-PanB interface is strikingly minimal and distributed across discontinuous regions, consistent with the weak affinity previously reported for this interaction (*14, 22, 23, 30*). This explains the difficulty of predicting pupylation sites from primary sequence or local structural features (*17, 22, 32, 34–36*). Notably, estimates of binding energies derived from the individual quaternary models predict affinities in the ∼100 µM to millimolar range (*37*), substantially weaker than the experimentally measured affinity (∼20 µM) (*14*). This discrepancy suggests that no single structural state is sufficient to account for the observed binding and instead supports a model in which affinity emerges from an ensemble of closely related, weakly interacting conformations.

### Continuous conformational variability underlies PafA target protein engagement

To determine whether substrate engagement occurs through discrete states or a continuum, we performed three-dimensional variability analysis (3DVA) on the consensus map (fig. S4), which revealed three distinct modes of continuous motion (fig. S12, movies S2-4). The first two components are characterized by differences in target protein engagement, with PafA adopting distinct orientations relative to PanB, suggesting multiple conformational states within the particle population. Component 1 is characterized by a rocking motion, whereby PafA transitions between peripheral and more central positions on the apical surface of PanB (fig. S12A, movie S2). Whereas component 2 captures a rotational motion, where PafA twists about the C5 symmetry axis of PanB (fig. S12B, movie S3). In contrast, the third component reflects modest, coordinated movements of both PafA and PanB (fig. S12C, movie S4). These results indicate that PafA engages substrates through a continuum of conformations rather than a few defined binding modes, consistent with a mechanism based on distributed, transient interactions (*38*).

### Substrate determinants of pupylation revealed by structure-guided mutagenesis of PanB

Previous studies have identified several PafA residues that are critical for its catalytic activity (*20–22,33*), and our cryoEM structure of the quaternary complex provides the first direct visualization of these residues in a structural context, explaining the effects of prior substitutions and demonstrating their mechanistic role in target protein engagement. For example, R201A and R207A substitutions in the extended loop nearly abolish pupylation activity (*21, 22*). In our model, PafA R201 forms a salt bridge with PanB E190, whereas R207 makes only limited contacts with PanB T214. Consistent with a broadly distributed and substrate-dependent interaction network, the contribution of individual residues likely varies across different substrates rather than reflecting universally conserved contacts. Additionally, a R442A substitution was shown to moderately reduce pupylation efficiency (*22*). Consistent with this, we show R442 interacts with the backbone carbonyl of PanB D32 in two of our models, providing a structural rationale for the previously observed functional effect.

To explore the complementary side of the interface, we leveraged our structural insights to design PanB variants targeting residues that interact with regions I and II of PafA, focusing on E190, T207, Q208, and T214 (Fig. 3G). All alanine-substituted PanB variants were soluble, and size-exclusion chromatography confirmed proper folding and oligomerization. PanB E190A exhibited the strongest reduction in pupylation, proceeding at only 0.17 ± 0.03 of the rate observed for wildtype PanB. E190 is contacted by multiple residues in PafA’s extended loop, highlighting its central role in substrate recognition. PanB Q208A, which loses contact with the PafA β3/4 loop was pupylated at 0.35 ± 0.04 of wildtype, while the PanB T214A variant, interacting with PafA R207 in the extended loop (4.9 Å away), decreased to 0.33 ± 0.03. Our quaternary complex models also reveal a previously unrecognized role for the PafA hinge in substrate engagement. The monomerized PafA_hinge_ variant showed reduced pupylation compared with wildtype PafA_monomer_ (Fig. 1). To probe this region further, we substituted PanB T207, which forms weak packing contacts with the hinge. The PanB T207A variant was pupylated at 0.28 ± 0.04 of the wildtype rate, demonstrating that even distal, weak interactions with the hinge contribute to substrate engagement. Together, these results show that multiple PanB residues, including those forming both central and distal contacts, contribute to efficient PafA-mediated pupylation, indicating that substrate recognition arises from the cumulative contribution of multiple weak interactions rather than a single dominant contact.

## Discussion

Substrate selection in the PPS has remained incompletely defined, partly due to the absence of a structure of PafA bound to a target protein. Here we present the first structures of PafA in complex with a model substrate. Our data support a mechanism in which PafA achieves broad substrate specificity through a sparse and distributed interaction surface, where affinity emerges from an ensemble of closely related conformations rather than a single defined complex. The minimal buried surface area observed in individual structural states, together with the discrepancy between predicted and measured binding affinities, indicates that substrate engagement is inherently dynamic. This behaviour is reminiscent of ‘fuzzy’ interactions (*39*), but here arises from a combination of structured contacts and flexible elements on both PafA and the substrate. Covalent tethering of Pup likely stabilizes these otherwise transient encounter states, biasing the ensemble toward catalytically productive configurations.

Consistent with this model, regions I (^50^SS^51^ hinge and β3/4 loop) and II (extended loop) are strictly conserved across PafA homologs (fig. S13). In contrast, residue R442 (region III) is not strictly conserved, but a positive charge is maintained in most species, suggesting that electrostatic interactions at this position are functionally important despite sequence variability (fig. S13). Across our four PafA-ADP-PanB∼Pup quaternary complex models, we observe variability in PafA-PanB interactions, which likely represent a subset of conformations that are sufficiently populated and structurally ordered to be resolved and modeled. Electrostatic hotspot residues R137, R218, and K444 (*22*), and extended loop residues A196, N205, D208 and E209 (*21*), previously identified as functionally important in PafA do not make contacts with PanB in our four models, which may reflect contributions from additional transient or heterogeneous conformations not captured in our data. In addition, a disordered loop from a neighbouring PanB subunit consistently approaches the PafA surface but lacks well-resolved density, suggesting that it may contribute transient contacts that are not captured in the structural models. These observations support a model in which PafA presents an extended interaction surface comprising multiple potential contact points, only a subset of which are engaged by any given substrate. In this framework, substrate recognition is governed by geometric compatibility rather than fixed sequence determinants, explaining the absence of consensus pupylation motifs (*17, 18, 22*). Although our analysis focuses on PanB, this mechanism likely generalizes across the pupylome and is consistent with the observed preference for high-molecular-weight substrates (*40*), which can engage multiple weak interactions simultaneously to achieve sufficient binding affinity.

The insights into substrate recognition by PafA also provide a framework for understanding its structural homolog, Dop, which catalyzes target protein depupylation (*20, 41–43*). In contrast to PafA, Dop recognizes the invariant Pup tag present on all substrates; however, given that pupylated proteins span a wide range of sizes and folds, Dop must likewise accommodate substantial substrate diversity beyond the Pup moiety. A distinguishing structural feature of Dop is the presence of the Dop loop, which is absent in PafA (*13, 20, 41*). Alignment of Dop [PDB: 7OXV (*13*)] with PafA in our quaternary complex model places this loop in a position equivalent to the pupylated PanB α8 helix, suggesting potential steric interference with substrate binding (fig. S14). While prior studies have proposed that this loop could act as either a positive or negative regulator of depupylation (*13, 41, 42, 44, 45*), our structural data support a model in which it functions primarily as a negative regulator based on the binding mode of PanB observed here. Together, these observations suggest that substrate recognition across the PPS is not dictated by primary sequence motifs, but instead relies on structural features, with PafA establishing specificity during pupylation and Dop operating through Pup-centric recognition while tolerating diverse protein structures. Definitive mechanistic understanding of Dop will require a structure of Dop bound to a pupylated substrate. Overall, these findings establish a structural framework for target protein recognition by PafA, helping to explain how it engages diverse targets, while also providing clues into how PafA and Dop shape the pupylome and proteasome-mediated protein degradation in prokaryotes.

## Acknowledgements

A.P. acknowledges support from a Natural Sciences and Engineering Research Council of Canada Postgraduate Doctoral Scholarship. A.B.U acknowledges support from the Fonds de Recherche du Québec-Santé (FRQ-S) doctoral fellowship. T.J.B.F acknowledges support from the Canadian Institutes of Health Research for a postdoctoral fellowship (FPE-194084). The Centre de Recherche en Biologie Structurale (CRBS) is funded by Fonds de Recherche du Québec (Health Sector) Research Centres Grant #288558. We thank K. Basu, and K. Sears at the Facility for Electron Microscopy Research (FEMR) at the McGill University for their assistance with microscope operation and data collection. FEMR is supported by the Canadian Foundation for Innovation, Quebec Government, and McGill University. Financial support was provided by a Canadian Institutes of Health Research Project Grant (PJT451412) to S.V., and Natural Sciences and Engineering Research Council of Canada Discovery Grants (RGPIN/03031-2022) to N.Z, and (RGPIN-2021-02843) to S.V. We thank Prof. Matthew Kimber (University of Guelph), Dr. Algirdas Velyvis (University of Guelph) for helpful discussions and critical review of the manuscript

## Data availability

Electron microscopy density maps have been deposited in the Electron Microscopy Databank (accession nos. EMD-76354, EMD-76355, EMD-76356, EMD-76357, EMD-76358, EMD-76359, EMD-76360, EMD-76361, EMD-76362, EMD-76363, EMD-76364, EMD-76365, EMD-76366, EMD-76367, EMD-76368, EMD-76369). Atomic models have been deposited in the Protein Databank (PDB ID 12DX, 12DY, 12DZ, 12EA).

## References

1. J. D. Etlinger, A. L. Goldberg, A soluble ATP-dependent proteolytic system responsible for the degradation of abnormal proteins in reticulocytes. Proc. Natl. Acad. Sci. U.S.A. 74, 54– 58 (1977).

2. G. Goldstein, M. Scheid, U. Hammerling, D. H. Schlesinger, H. D. Niall, E. A. Boyse, Isolation of a polypeptide that has lymphocyte-differentiating properties and is probably represented universally in living cells. Proc. Natl. Acad. Sci. U.S.A. 72, 11–15 (1975).

3. A. L. Haas, J. V. Warms, A. Hershko, I. A. Rose, Ubiquitin-activating enzyme: Mechanism and role in protein-ubiquitin conjugation. J. Biol. Chem. 257, 2543–2548 (1982).

4. A. L. Goldberg, Protein degradation and protection against misfolded or damaged proteins. Nature 426, 895–899 (2003).

5. K. E. Burns, K. H. Darwin, Pupylation: A signal for proteasomal degradation in Mycobacterium tuberculosis. Subcell. Biochem. 54, 149–157 (2010).

6. K. H. Darwin, S. Ehrt, J.-C. Gutierrez-Ramos, N. Weich, C. F. Nathan, The proteasome of Mycobacterium tuberculosis is required for resistance to nitric oxide. Science 302, 1963– 1966 (2003).

7. F. Striebel, F. Imkamp, D. Özcelik, E. Weber-Ban, Pupylation as a signal for proteasomal degradation in bacteria. Biochem. Biophys. Acta 1843, 103–113 (2014).

8. M. I. Samanovic, H. Li, K. H. Darwin, The Pup-proteasome system of Mycobacterium tuberculosis. Subcell. Biochem. 66, 267–295 (2013).

9. M. I. Samanovic, S. Tu, O. Novák, L. M. Iyer, F. E. McAllister, L. Aravind, S. P. Gygi, S. R. Hubbard, M. Strnad, K. H. Darwin, proteasomal control of cytokinin synthesis protects Mycobacterium tuberculosis against nitric oxide. Mol. Cell. 57, 984–994 (2015).

10. M. Sutter, F. F. Damberger, F. Imkamp, F. H.-T. Allain, E. Weber-Ban, Prokaryotic ubiquitin-like protein (pup) is coupled to substrates via the side chain of its c-terminal glutamate. J. Am. Chem. Soc. 132, 5610–5612 (2010).

11. F. Striebel, F. Imkamp, M. Sutter, M. Steiner, A. Mamedov, E. Weber-Ban, Bacterial ubiquitin-like modifier Pup is deamidated and conjugated to substrates by distinct but homologous enzymes. Nat. Struct. Mol. Biol. 16, 647–651 (2009).

12. M. Bolten, C. Vahlensieck, C. Lipp, M. Leibundgut, N. Ban, E. Weber-Ban, Depupylase dop requires inorganic phosphate in the active site for catalysis. J. Biol. Chem. 292, 4044– 4053 (2017).

13. H. Cui, A. U. Müller, M. Leibundgut, J. Tian, N. Ban, E. Weber-Ban, Structures of prokaryotic ubiquitin-like protein Pup in complex with depupylase Dop reveal the mechanism of catalytic phosphate formation. Nat. Commun. 12, 6635 (2021).

14. E. Guth, M. Thommen, E. Weber-Ban, Mycobacterial ubiquitin-like protein ligase PafA follows a two-step reaction pathway with a phosphorylated Pup intermediate. J. Biol. Chem. 286, 4412–4419 (2011).

15. J. Laederach, H. Cui, E. Weber-Ban, Pupylated proteins are subject to broad proteasomal degradation specificity and differential depupylation. PLoS ONE 14, e0215439 (2019).

16. J. Barandun, C. L. Delley, N. Ban, E. Weber-Ban, Crystal structure of the complex between prokaryotic ubiquitin-like protein and its ligase PafA. J. Am. Chem. Soc. 135, 6794–6797 (2013).

17. J. Watrous, K. Burns, W.-T. Liu, A. Patel, V. Hook, V. Bafna, C. E. Barry, S. Bark, P. C. Dorrestein, Expansion of the mycobacterial “PUPylome.” Mol. BioSyst. 6, 376–385 (2010).

18. X. Chen, J.-D. Qiu, S.-P. Shi, S.-B. Suo, R.-P. Liang, Systematic analysis and prediction of pupylation sites in prokaryotic proteins. PLoS ONE 8, e74002 (2013).

19. Q. Liu, J. Zheng, W. Sun, Y. Huo, L. Zhang, P. Hao, H. Wang, M. Zhuang, A proximity-tagging system to identify membrane protein–protein interactions. Nat. Methods 15, 715– 722 (2018).

20. D. Özcelik, J. Barandun, N. Schmitz, M. Sutter, E. Guth, F. F. Damberger, F. H.-T. Allain, N. Ban, E. Weber-Ban, Structures of Pup ligase PafA and depupylase Dop from the prokaryotic ubiquitin-like modification pathway. Nat. Commun. 3, 1014 (2012).

21. O. Regev, M. Korman, N. Hecht, Z. Roth, N. Forer, R. Zarivach, E. Gur, An extended loop of the Pup ligase, PafA, mediates interaction with protein targets. J. Mol. Biol. 428, 4143– 4153 (2016).

22. M. F. Block, C. L. Delley, L. M. L. Keller, T. T. Stuehlinger, E. Weber-Ban, Electrostatic interactions guide substrate recognition of the prokaryotic ubiquitin-like protein ligase PafA. Nat. Commun. 14, 5266 (2023).

23. E. Krissinel, K. Henrick, Inference of macromolecular assemblies from crystalline state. J. Mol. Biol. 372, 774–797 (2007).

24. S. C. Kahne, J. H. Yoo, J. Chen, K. Nakedi, L. M. Iyer, G. Putzel, N. M. Samhadaneh, A. Pironti, L. Aravind, D. C. Ekiert, G. Bhabha, K. Y. Rhee, K. H. Darwin, Identification of a depupylation regulator for an essential enzyme in Mycobacterium tuberculosis. Proc. Natl. Acad. Sci. U.S.A. 121, e2407239121 (2024).

25. E. Krissinel, K. Henrick, Secondary-structure matching (SSM), a new tool for fast protein structure alignment in three dimensions. Acta Crystallogr. D Biol. Crystallogr. 60, 2256– 2268 (2004).

26. A. Plourde, J. C. Ogata-Bean, S. Vahidi, Mapping the structural heterogeneity of Pup ligase PafA using H/D exchange mass spectrometry. J. Biol. Chem. 301, 108437 (2025).

27. M. J. Pearce, P. Arora, R. A. Festa, S. M. Butler-Wu, R. S. Gokhale, K. H. Darwin, Identification of substrates of the Mycobacterium tuberculosis proteasome. EMBO J. 25, 5423–5432 (2006).

28. B. N. Chaudhuri, M. R. Sawaya, C.-Y. Kim, G. S. Waldo, M. S. Park, T. C. Terwilliger, T.O. Yeates, the crystal structure of the first enzyme in the pantothenate biosynthetic pathway, ketopantoate hydroxymethyltransferase, from M. tuberculosis. Structure 11, 753–764 (2003).

29. F. Von Delft, T. Inoue, S. A. Saldanha, H. H. Ottenhof, F. Schmitzberger, L. M. Birch, V. Dhanaraj, M. Witty, A. G. Smith, T. L. Blundell, C. Abell, Structure of E. coli ketopantoate hydroxymethyl transferase complexed with ketopantoate and mg2+, solved by locating 160 selenomethionine sites. Structure 11, 985–996 (2003).

30. N. Ofer, N. Forer, M. Korman, M. Vishkautzan, I. Khalaila, E. Gur, Allosteric transitions direct protein tagging by PafA, the prokaryotic ubiquitin-like protein (Pup) ligase. J. Biol. Chem. 288, 11287–11293 (2013).

31. J. Barandun, F. F. Damberger, C. L. Delley, J. Laederach, F. H. T. Allain, E. Weber-Ban, Prokaryotic ubiquitin-like protein remains intrinsically disordered when covalently attached to proteasomal target proteins. BMC Struct. Biol. 17, 1 (2018).

32. Md. M. Hasan, Y. Zhou, X. Lu, J. Li, J. Song, Z. Zhang, Computational identification of protein pupylation sites by using profile-based composition of k-spaced amino acid pairs. PLoS ONE 10, e0129635 (2015).

33. F. A. Cerda-Maira, M. J. Pearce, M. Fuortes, W. R. Bishai, S. R. Hubbard, K. H. Darwin, Molecular analysis of the prokaryotic ubiquitin-like protein (Pup) conjugation pathway in Mycobacterium tuberculosis: Pup conjugation pathway. Mol. Microbiol. 77, 1123–1135 (2010).

34. F. A. Cerda-Maira, F. McAllister, N. J. Bode, K. E. Burns, S. P. Gygi, K. H. Darwin, Reconstitution of the Mycobacterium tuberculosis pupylation pathway in Escherichia coli. EMBO Rep. 12, 863–870 (2011).

35. V. Singh, A. Sharma, A. Dehzangi, T. Tsunoda, PupStruct: prediction of pupylated lysine residues using structural properties of amino acids. Genes 11, 1431 (2020).

36. X. Nan, L. Bao, X. Zhao, X. Zhao, A. Sangaiah, G.-G. Wang, Z. Ma, EPuL: An enhanced positive-unlabeled learning algorithm for the prediction of pupylation sites. Molecules 22, 1463 (2017).

37. R. V. Honorato, P. I. Koukos, B. Jiménez-García, A. Tsaregorodtsev, M. Verlato, A. Giachetti, A. Rosato, A. M. J. J. Bonvin, Structural biology in the clouds: the WeNMR-EOSC ecosystem. Front. Mol. Biosci. 8, 729513 (2021).

38. S. C. Kahne, K. H. Darwin, “Pupdates” on proteasomal degradation in bacteria. J. Bacteriol. 207, e00111–25 (2025).

39. M. Fuxreiter, Fuzziness in protein interactions—a historical perspective. J. Mol. Biol. 430, 2278–2287 (2018).

40. Y. Elharar, Z. Roth, N. Hecht, R. Rotkopf, I. Khalaila, E. Gur, Posttranslational regulation of coordinated enzyme activities in the Pup-proteasome system. Proc. Natl. Acad. Sci. U.S.A. 113, E1605–E1614 (2016).

41. J. H. Yoo, S. C. Kahne, K. H. Darwin, A conserved loop sequence of the proteasome system depupylase Dop regulates substrate selectivity in Mycobacterium tuberculosis. J. Biol. Chem. 298, 102478 (2022).

42. F. Imkamp, T. Rosenberger, F. Striebel, P. M. Keller, B. Amstutz, P. Sander, E. Weber-Ban, Deletion of dop in Mycobacterium smegmatis abolishes pupylation of protein substrates in vivo. Mol. Microbiol. 75, 744–754 (2010).

43. F. Imkamp, F. Striebel, M. Sutter, D. Özcelik, N. Zimmermann, P. Sander, E. Weber-Ban, Dop functions as a depupylase in the prokaryotic ubiquitin-like modification pathway. EMBO Rep. 11, 791–797 (2010).

44. N. Hecht, C. L. Monteil, G. Perrière, M. Vishkautzan, E. Gur, Exploring Protein Space: From Hydrolase to Ligase by Substitution. Mol. Biol. Evol. 38, 761–776 (2021).

45. N. Hecht, M. Becher, M. Korman, M. Vishkautzan, E. Gur, Inter- and intramolecular regulation of protein depupylation in Mycobacterium smegmatis. FEBS J. 287, 4389–4400 (2020).

46. M. Schorb, I. Haberbosch, W. J. H. Hagen, Y. Schwab, D. N. Mastronarde, Software tools for automated transmission electron microscopy. Nat. Methods 16, 471–477 (2019).

47. A. Punjani, J. L. Rubinstein, D. J. Fleet, M. A. Brubaker, cryoSPARC: algorithms for rapid unsupervised cryo-EM structure determination. Nat. Methods 14, 290–296 (2017).

48. T. Bepler, A. Morin, M. Rapp, J. Brasch, L. Shapiro, A. J. Noble, B. Berger, Positive-unlabeled convolutional neural networks for particle picking in cryo-electron micrographs. Nat. Methods 16, 1153–1160 (2019).

49. J. Zivanov, T. Nakane, S. H. W. Scheres, Estimation of high-order aberrations and anisotropic magnification from cryo-EM data sets in RELION-3.1. IUCrJ 7, 253–267 (2020).

50. S. H. W. Scheres, RELION: Implementation of a Bayesian approach to cryo-EM structure determination. J. Struct. Biol. 180, 519–530 (2012).

51. A. Punjani, D. J. Fleet, 3D variability analysis: Resolving continuous flexibility and discrete heterogeneity from single particle cryo-EM. J. Struct. Biol. 213, 107702 (2021).

52. D. Liebschner, P. V. Afonine, M. L. Baker, G. Bunkóczi, V. B. Chen, T. I. Croll, B. Hintze, L.-W. Hung, S. Jain, A. J. McCoy, N. W. Moriarty, R. D. Oeffner, B. K. Poon, M. G. Prisant, R. J. Read, J. S. Richardson, D. C. Richardson, M. D. Sammito, O. V. Sobolev, D. H. Stockwell, T. C. Terwilliger, A. G. Urzhumtsev, L. L. Videau, C. J. Williams, P. D. Adams, Macromolecular structure determination using X-rays, neutrons and electrons: recent developments in Phenix. Acta Crystallogr. D Struct. Biol. 75, 861–877 (2019).

53. E. C. Meng, T. D. Goddard, E. F. Pettersen, G. S. Couch, Z. J. Pearson, J. H. Morris, T. E. Ferrin, UCSF CHIMERAX: Tools for structure building and analysis. Protein Sci. 32, e4792 (2023).

54. J. Abramson, J. Adler, J. Dunger, R. Evans, T. Green, A. Pritzel, O. Ronneberger, L. Willmore, A. J. Ballard, J. Bambrick, S. W. Bodenstein, D. A. Evans, C.-C. Hung, M. O’Neill, D. Reiman, K. Tunyasuvunakool, Z. Wu, A. Žemgulytė, E. Arvaniti, C. Beattie, O. Bertolli, A. Bridgland, A. Cherepanov, M. Congreve, A. I. Cowen-Rivers, A. Cowie, M. Figurnov, F. B. Fuchs, H. Gladman, R. Jain, Y. A. Khan, C. M. R. Low, K. Perlin, A. Potapenko, P. Savy, S. Singh, A. Stecula, A. Thillaisundaram, C. Tong, S. Yakneen, E. D. Zhong, M. Zielinski, A. Žídek, V. Bapst, P. Kohli, M. Jaderberg, D. Hassabis, J. M. Jumper, Accurate structure prediction of biomolecular interactions with AlphaFold 3. Nature 630, 493–500 (2024).

55. F. Long, R. A. Nicholls, P. Emsley, S. Gražulis, A. Merkys, A. Vaitkus, G. N. Murshudov, AceDRG : a stereochemical description generator for ligands. Acta Crystallogr. D Struct. Biol. 73, 112–122 (2017).

56. P. Emsley, B. Lohkamp, W. G. Scott, K. Cowtan, Features and development of Coot. Acta Crystallogr. D Biol. Crystallogr. 66, 486–501 (2010).

57. I. Hutařová Vařeková, J. Hutař, A. Midlik, V. Horský, E. Hladká, R. Svobodová, K. Berka, 2DProts: database of family-wide protein secondary structure diagrams. Bioinformatics 37, 4599–4601 (2021).

