## Supplemental Material for "Structural basis of substrate recognition for proteasome degradation by prokaryotic ubiquitin-like protein ligase PafA"

**This PDF includes:**

Materials and methods

Figs. S1-14

Caption for Table S1

Table S2

Caption for Movies S1-S4

References

**Other Supplementary Material for this manuscript include:**

Table S1 (.xlsx)

Movies S1-S4 (.mp4)

### Materials and Methods

*Plasmids, protein production and protein purification*

All genes were codon-optimized for expression in *Escherichia coli* in pET24, pEt28a, or pET29a vectors, and encode an N-terminal $His_6$-SUMO tag for protein purification. Plasmids encoding wildtype *Corynebacterium glutamicum* PafA, *C. glutamicum* $Pup^E$, and wildtype *Mycobacterium tuberculosis* PanB were obtained from BioBasic Canada. Plasmids for *C. glutamicum* PafA variants and *M. tuberculosis* PanB variants were obtained from Twist Bioscience.

The plasmids were transformed into T7 Express *lysY* competent *E. coli* (New England Biolabs). The cells were grown in LB broth supplemented with 100 mg/L $MgCl_2$ and 30 mg/L kanamycin, at 37 °C with 180 rpm agitation to an optical density at 600 nm of 0.6 to 0.8. Cells were induced with 0.2 mM isopropyl β-D-1-thiogalactopyranoside (IPTG) to overproduce proteins at 30 °C for 18-20 hours for PafA constructs, 18 °C for 18-20 hours for PanB constructs, or 37 °C for 4 hours for $Pup^E$. Following protein overproduction, cells were harvested by centrifugation at 5000 × *g* relative centrifugal force (RCF) for 30 min at 4 °C. The cell pellets were resuspended in Buffer A (50 mM Tris-HCl (pH 8.0), 300 mM KCl, 20 mM imidazole) and either frozen at -20 °C or immediately used for purification.

Cells were lysed via homogenization in an EmulsiFlex-C3 (Avestin) at 1000 bar and the supernatant was cleared via centrifugation at 30000-45000 × *g* RCF for 30 minutes at 4 °C. The soluble fraction was loaded onto an immobilized metal affinity chromatography (IMAC) column with chelating sepharose beads (Cytiva) charged with $Ni^{2+}$. Column washes were performed with Buffer A and Buffer A supplemented with 50 mM imidazole. All proteins were eluted via Buffer A supplemented with 500 mM imidazole. PafA and PanB constructs were incubated with Ubiquitin-like specific protease 1 (Ulp1) overnight at 4 °C to remove the His-SUMO tag, while being dialyzed against buffer A supplemented with 1 mM dithiothreitol (DTT). PafA and PanB were subjected to a second round of IMAC purification in which the flowthrough, containing mature PafA or PanB, was retained. Proteins were concentrated using centrifugal concentrators (Amicon) with molecular weight cut-offs (MWCO) of 50 kDa for PafA and PanB, or 10 kDa for $His_6$-SUMO-$Pup^E$ to 0.5 mL. The concentrated proteins were applied to a Superdex 200 Increase 10/300 GL (Cytiva) equilibrated in 20 mM HEPES (pH 7.5) with 150 mM NaCl at either 4 °C or room temperature (20-25 °C). Fractions were collected and sample purity was evaluated using Coomassie staining and SDS-PAGE analysis. $His_6$-SUMO-$Pup^E$ and PanB constructs were either stored at 4 °C or supplemented with 10% [v/v] glycerol, flash frozen in liquid nitrogen, and stored at -80 °C. PafA constructs were stored at 4 °C. In experiments involving wildtype $PafA_{monomer}$, the protein was used immediately following size-exclusion chromatography separation from wildtype $PafA_{dimer}$.

*Analytical size-exclusion chromatography*

10 µM of wildtype PafA and PafA variants were applied to an ACQUITY Premier Protein 250 Å column (Waters) on an Agilent 1100 series high performance liquid chromatography (HPLC) instrument equilibrated in 20 mM HEPES (pH 7.5) with 150 mM NaCl. UV detection at 220 nm was recorded with a G1314A variable wavelength detector (Agilent). Samples were injected from a G1367 autosampler (Agilent) at ambient room temperature.

*in vitro reconstitution of pupylation system*

$His_6$-SUMO-$Pup^E$ was incubated with Ulp1 and 1 mM DTT for 18-20 hours at 4 °C. For activity profiling, the pupylation system was reconstituted *in vitro* at 100 µL with 3 µM PafA, 10 µM PanB (wildtype and variants), 20 µM cleaved $Pup^E$, and 3 mM ATP, in buffer containing 20 mM HEPES (pH 7.5), 150 mM NaCl, and 20 mM $MgCl_2$. The pupylation reaction proceeded at room temperature (20-25 °C) and was quenched with equal volume of Laemmli buffer. The samples were separated using SDS-PAGE and stained with Coomassie. Densitometry analysis was performed using ImageJ 0.6.0 (National Institute of Health). Baseline density was removed using rolling ball background subtraction. Density plots for rows containing the substrate and product bands were obtained using the Gel Analyzer tool. Plots were manually divided into lanes and band densities were measured as the area under the curve for each lane. The ratio between product band density and the substrate band density at the reaction onset was determined and normalized to 1. These data were fit to an exponential model where the asymptote was fixed at 1 and the rate constant was allowed to float. Monte Carlo simulation with 1000 iterations was performed to determine the uncertainties in the fitted rate constant. These data were fit to an exponential model where the asymptote was fixed at 1 and the rate constant was allowed to float. Monte Carlo simulation with 1000 iterations was performed to determine the uncertainties in the fitted rate constant.

For the generation of high yields of PanB~Pup product, a 10 mL reaction was prepared in 20 mM HEPES (pH 7.5), 150 mM NaCl, 20 mM $MgCl_2$, and 1 mM DTT. 1 µM of $PafA_{R447G}$ was incubated with 80 µM PanB and 80 µM $His_6$-SUMO-$Pup^E$ and Ulp1. $His_6$-SUMO-$Pup^E$ cleavage (with Ulp1) proceeded at the same time as PanB pupylation. The reaction proceeded for 30 hours at 4 °C and was not quenched upon completion.

*Isolation of PanB~Pup and the quaternary complex*

The 10 mL pupylation reaction was concentrated by centrifugation with a 50 kDa MWCO concentrator (Amicon) to 0.5 mL. The concentrated protein was applied to a 200 Increase 10/300 GL SEC column (Cytiva) equilibrated in 20 mM HEPES (pH 7.5) with 150 mM NaCl at 4 °C. Fractions containing pupylated PanB (PanB~Pup), as assessed by SDS-PAGE analysis, were pooled. Band densitometry measurements with ImageJ 0.6.0 (National Institute of Health) were used to determine the fraction of pupylated PanB subunits. The concentration of PanB~Pup was determined using the 280 nm extinction coefficient of PanB only, as Pup has an $\varepsilon$ of 0.

200 µM PanB~Pup (PanB subunit concentration) was incubated with 175 µM $PafA_{D64N/R447G}$ for one hour at room temperature (20-25 °C) in 20 mM HEPES (pH 7.5), 150 mM NaCl, 0.5 mM ATP, and 1 mM $MgCl_2$ to promote complex assembly. This was applied to a Superdex 200 Increase 10/300 GL SEC column (Cytiva) equilibrated in the same buffer to isolate the quaternary complex from unbound $PafA_{D64N/R447G}$. SEC fractions containing the quaternary complex, as assessed by SDS-PAGE analysis, were pooled and concentrated to 17 mg/mL with a 50 kDa MWCO concentrator (Amicon) and stored at 4 °C.

*Electron cryomicroscopy sample preparation*

IGEPAL CA-630 was added to the 17 mg/mL quaternary complex sample at a final concentration of 0.025% to ensure heterogenous particle orientation during vitrification. The sample was then applied to a holey carbon grid (C-flat CF-1.2/1.3-3Cu-T) that had been glow discharged in air at 5 mA for 15 s using a Pelco easiGlow Glow Discharge Cleaning System. Sample vitrification was performed in a Vitrobot Mark IV (Thermo Fisher Scientific) at 4 °C and 100% humidity. The sample was incubated on the grid for 3 s and the grid was blotted for 4 s with a blot force of 1 before plunging in liquid ethane.

*Electron cryomicroscopy data collection*

Datasets were collected using the SerialEM software (*46*) on a Titan Krios microscope (Thermo Fisher Scientific). Movies were recorded on a Gatan K3 direct electron detector equipped with a Quantum LS imaging filter. The total electron dose per movie was 60 $e^-/Å^2$, equally spread over 40 frames. Data were collected at a magnification of x130,000, yielding images with a calibrated pixel size of 0.675 Å. The nominal defocus range used during data collection was between -1.00 µm and -2.5 µm. A summary of the data collection parameters for all datasets can be found in Table S1.

*Electron cryomicroscopy data processing*

All cryo-EM data processing was performed using CryoSPARCv4 (*47*) . Movie frames were aligned and corrected for beam-induced motion using Patch Motion Correction and contrast transfer function (CTF) parameters were estimated using Patch CTF Estimation. Micrographs with CTF fit resolution worse than 10 Å, or with unusually high full-frame motion were discarded. The remaining micrographs were used for downstream processing.

Initial particle picking was performed using the Blob Picker with particle diameters set to 70–200 Å. Extracted particles were subjected to 2D classification, followed by *ab initio* reconstruction and heterogenous refinement. Classes corresponding to projections of the target complex were selected, and the resulting 2D class averages were used as templates for a second round of particle picking with the Template Picker. Template-based picked particles were processed in the same manner through 2D classification, *ab initio* reconstruction and heterogenous refinement. Particles assigned to classes consistent with the target complex were then used to train the Topaz particle picking neural network (*48*). The trained Topaz model was subsequently used to identify additional particles from the full dataset. These particles were again subjected to 2D classification, *ab initio* reconstruction and heterogenous refinement to enrich for particles corresponding to the target complex. To maximize particle recovery, particles retained from all three picking approaches were merged after duplicate particle removal and subjected to another round of *ab initio* reconstruction and heterogeneous refinement. Classes exhibiting prominent density for PafA were pooled and subjected to reference-based motion correction in CryoSPARC to perform per-particle motion correction. The motion-corrected particles were then used to generate a consensus map using non-uniform refinement combined with global and local CTF refinements (*49*). In parallel, these particles were exported to RELION (*50*) for 3D classification to further assess structural heterogeneity. Although this analysis identified classes containing more than one bound PafA molecule, these classes contained too few particles for further refinement. Particles from this stage were also used for 3D variability analysis (3DVA) (*51*) in CryoSPARC to probe conformational heterogeneity in PafA binding to PanB. 3DVA was performed using five components and a filter resolution of 4 Å.

To better resolve PafA and its binding interface with PanB, focused 3D classification was performed in CryoSPARC using the consensus map, and a mask encompassing the PafA-binding region(s). Of the ten classes, four displaying well-resolved secondary structure features for PafA were selected for separate Non-uniform refinements. Focussed local refinements on the PafA and PanB regions were then performed for each of these four classes to further improve map quality. The resulting focussed maps were combined in PHENIX (version 2.0_5936) (*52*) using the "combine focused maps" utility to generate composite maps for the four classes.

ChimeraX (version 1.9) (*53*) was used for map visualization and figure preparation.

*Model building and refinement*

The PanB decamer structure (PDB: 1OY0 (*28*)) and the AlphaFold3 (*54*) predicted model of the PafA-Pup complex were used as initial models. These were docked onto the composite maps using ChimeraX (*53*) to generate the starting model. Restraints for the isopeptide bond were generated using AceDRG (*55*) in Coot (*56*). Model building was performed by iterative cycles of manual rebuilding in Coot (*56*) and real-space refinement in PHENIX (*52*), resulting in improved model geometry and agreement with the density map. Final model validation was also performed on PHENIX (*52*), and refinement and validation statistics are summarized in Table S1.

**Supplemental table 1. Cryo-EM data collection, model building and refinement statistics.Supplemental movie 1. Morph between the four quaternary complex classes highlighting the PanB~Pup isopeptide linkage and the different interface regions.**

**Supplemental movie 2. 3D variability analysis Component 1 depicting PafA rocking.**

**Supplemental movie 3. 3D variability analysis Component 2 depicting PafA twisting.**

**Supplemental movie 4. 3D variability analysis Component 3 depicting coordinated PafA-PanB movement.**

**Supplemental table 2. Summary of PafA-PanB contacts within 5 Å.** Contacts between atoms in PafA and PanB within 5 Å are shown, categorized by PafA region. These regions include the $^{50}$SS$^{51}$ hinge and β3/4 loop (region I), the extended loop (region II), and CTD residue R442 (region III). Contacts are indicated by a checkmark (✓). Residues that have been mutated in previous studies are indicated by [†], and residues that have been mutated here are indicated by [*].

| PafA region | PafA residue | PanB residue | Quaternary complex class | | | |
|---|---|---|---|---|---|---|
| | | | 1 | 2 | 3 | 4 |
| Hinge and β3/4 loop (I) | S51 | E204 | ✓ | | ✓ | ✓ |
| | L63 | Q208[*] | ✓ | ✓ | ✓ | ✓ |
| | D64N[†] | Q208[*] | ✓ | ✓ | ✓ | ✓ |
| | | K212[†] | | ✓ | ✓ | |
| | V65[†] | I182 | ✓ | ✓ | ✓ | ✓ |
| | | Q208[*] | ✓ | ✓ | ✓ | ✓ |
| | | K212[†] | ✓ | ✓ | ✓ | ✓ |
| | G66 | Q208[*] | ✓ | ✓ | ✓ | ✓ |
| Extended loop (II) | T197[†] | E190 | ✓ | ✓ | | |
| | T198[†] | E136 | | ✓ | | |
| | | A187 | | ✓ | | |
| | | E190[*] | | ✓ | ✓ | |
| | R199[†] | T146 | ✓ | | | |
| | | I186 | | ✓ | | |
| | | A189 | ✓ | | | |
| | | E190[*] | ✓ | ✓ | ✓ | |
| | | A191 | ✓ | | | |
| | | G192 | ✓ | | | |
| | R201[†] | E190[*] | ✓ | | | |
| | R207[†] | T214[*] | ✓ | | | |
| | P210 | A31 | ✓ | | ✓ | |
| | | R32 | ✓ | | ✓ | |
| | H211[†] | A31 | ✓ | | | |
| | | R32 | ✓ | | ✓ | |
| | A212 | R32 | ✓ | | | |
| CTD (III) | R442[†] | A31 | | ✓ | | ✓ |
| | | R32 | | ✓ | | ✓ |
| | | G33 | | ✓ | | |

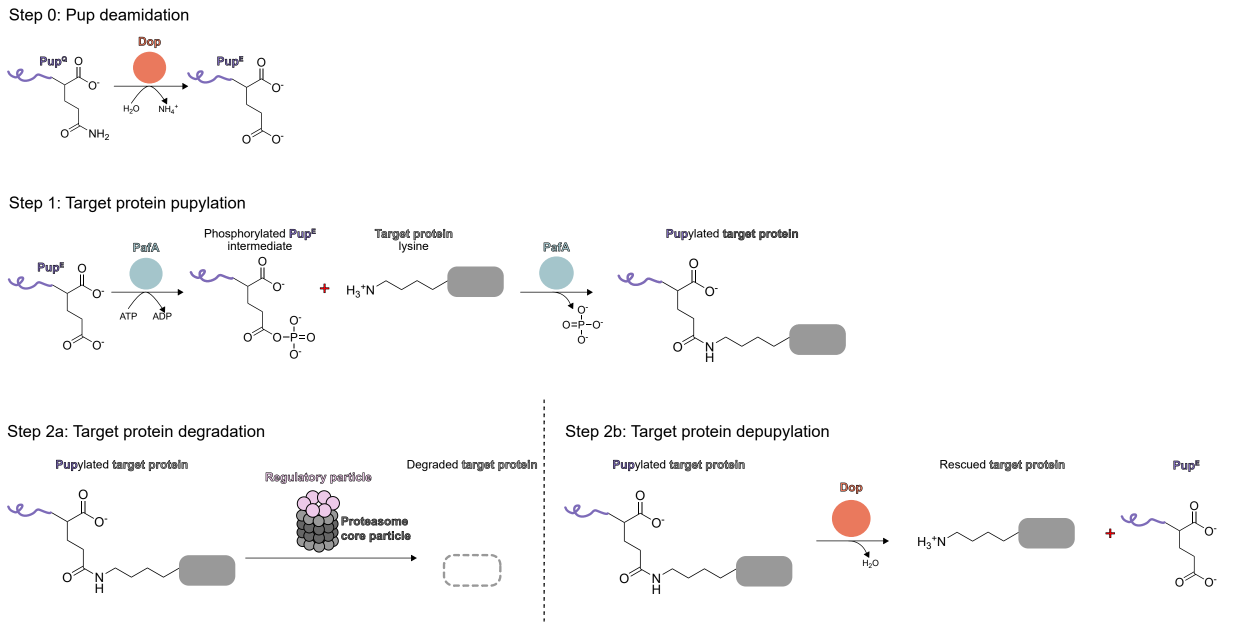


**Supplemental figure 1. Pup-proteasome system schematic**. Pup deamidation (step 0) is required in species that encode Pup$^Q$. Pup$^E$ is then ligated to a target protein in a two-step reaction catalyzed by PafA (step 1). The resulting pupylated target can either be degraded by the proteasome machinery (step 2a) or depupylated by Dop to rescue the protein (step 2b).

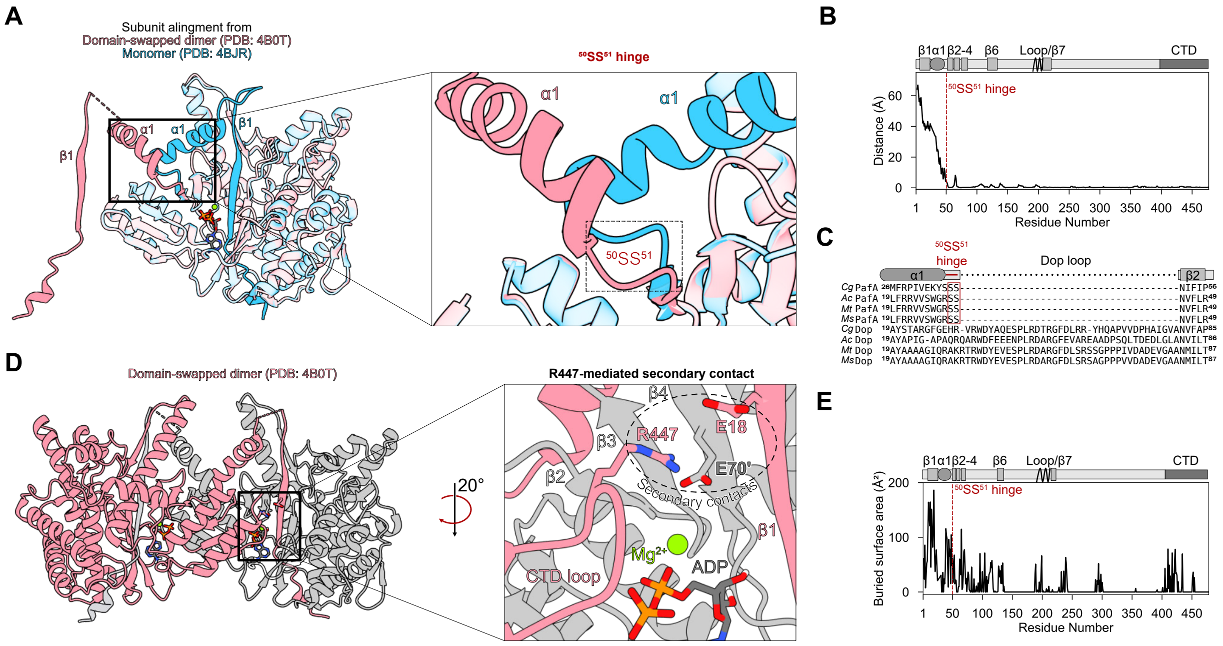


**Supplemental figure 2. Design of PafA monomerization variants. A.** Alignment of a PafA subunit from the domain-swapped dimer (PDB: 4B0T (*20*) - blue) and monomer [PBD: 4BJR (*16*) – pink] structures of PafA. The N-terminal strand-helix motif interacts with the other subunit in the domain-swapped dimer. The remainder of the structure aligns with an RMSD of 0.5 Å. **B.** One PafA subunit from the domain-swapped dimer [PDB: 4B0T (*20*) ] and monomer [PDB: 4BJR (*16*) ] were aligned and the distance between Cα were calculated. The N-terminal strand-helix motif (β1 and α1) differ greatly between the two structures. The distance between the Cα dramatically drop at the $^{50}$SS$^{51}$ hinge and remain closely aligned between the two subunits for the rest of the sequence. The N-terminal strand-helix motif (β1 and α1), remainder of the active site cradle (β2-4, β6, and β7), and the extended loop (residues 196-216) are indicated above the plot. **C.** Curated multiple sequence alignment of 65 PafA and Dop sequences from the UniprotKB database (*3*). PafA and Dop are structural homologs that likely arose from a gene duplication event and catalyze protein pupylation and depupylation, respectively (*1*, *4*). The alignment of sequences for Dop and PafA from *Mycobacterium tuberculosis* (*Mt*), *M. smegmatis* (*Ms*), *Corynebacterium glutamicum* (*Cg*), and *Acidothermus cellulolyticus* (*Ac*) are displayed. The conserved $^{50}$SS$^{51}$ hinge in PafA immediately precedes the roughly 40-amino acid conserved loop in Dop. **D.** The domain-swapped dimer [PDB: 4B0T (*20*) ] is shown, coloured by chain. The C-terminal domain (CTD) forms secondary contacts within the domain-swapped structure, importantly R447 interacts with the active site of the other subunit. **E.** Buried surface area (Å2) per residue in the domain-swapped $PafA_{dimer}$ [PDB: 4B0T (*20*) ]. The $PafA_{dimer}$ interface buries ~6300 Å2 surface area, involving the active site, extended loop, and C-terminal domain. The N-terminal strand-helix motif (β1 and α1), remainder of the active site cradle (β2-4, β6, and β7), and the extended loop (residues 196-216) are indicated above the plot.

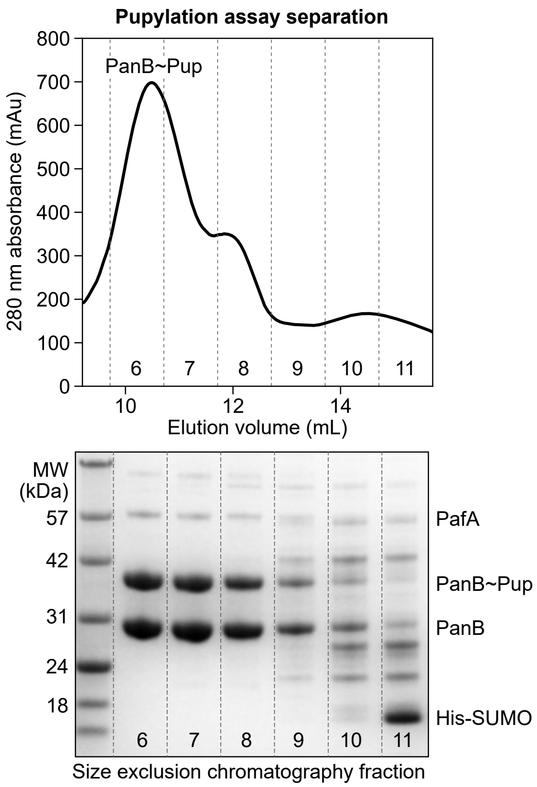


**Supplemental figure 3. PanB~Pup product isolation.** SEC profile and corresponding gel for the purification of PanB~Pup. PanB was pupylated (PanB~Pup) *in vitro*, followed by SEC-based isolation to remove lower molecular weight contaminants.

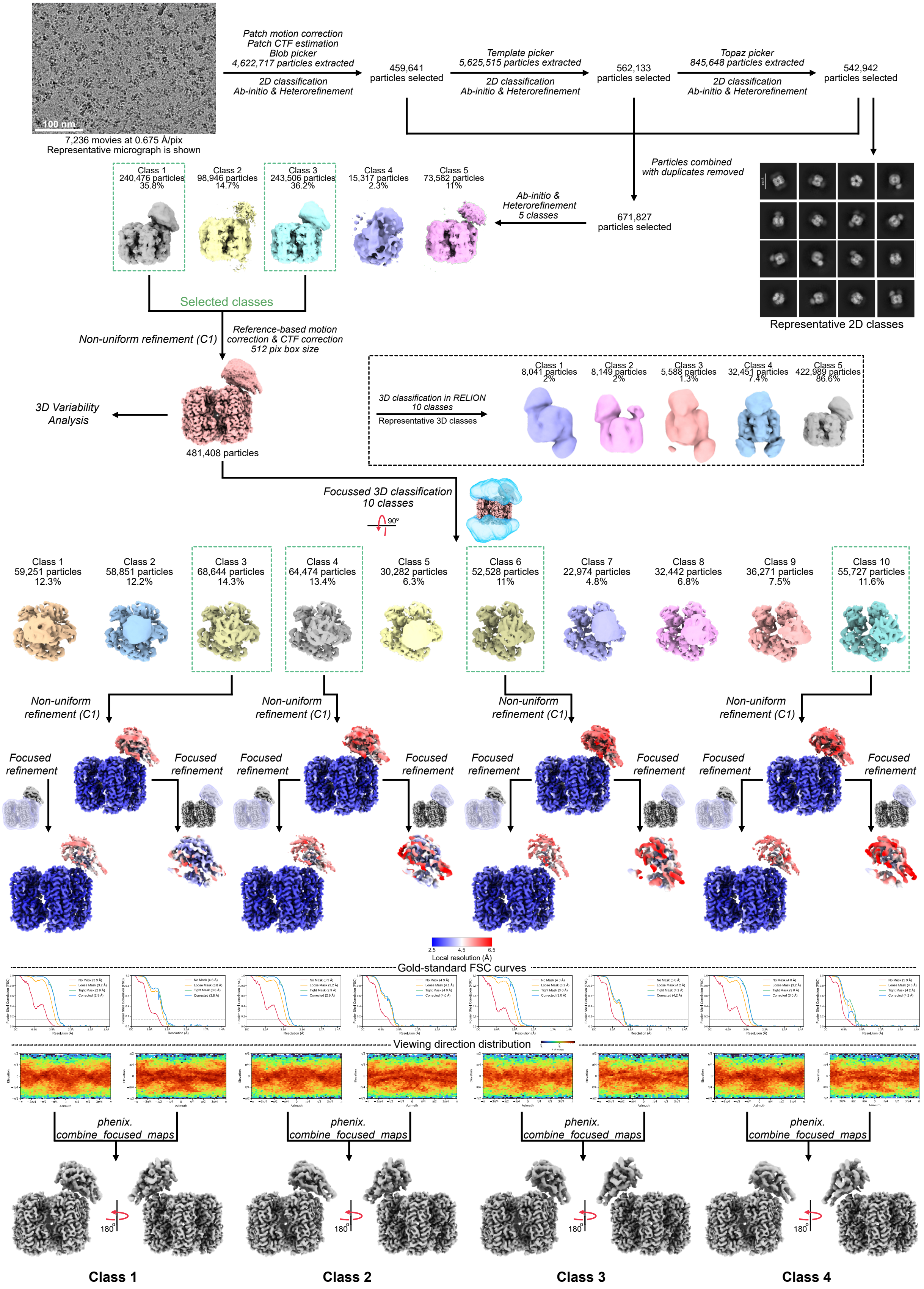

Patch motion correction
Patch CTF estimation
Blob picker
4,622,717 particles extracted
2D classification
Ab-initio & Heterorefinement
459,641 particles selected
Template picker
5,625,515 particles extracted
2D classification
Ab-initio & Heterorefinement
562,133 particles selected
Topaz picker
845,648 particles extracted
2D classification
Ab-initio & Heterorefinement
542,942 particles selected
100 nm
7,236 movies at 0.675 Å/pix
Representative micrograph is shown
Particles combined with duplicates removed
Ab-initio & Heterorefinement 5 classes
671,827 particles selected
Class 1 240,476 particles 35.8%
Class 2 98,946 particles 14.7%
Class 3 243,506 particles 36.2%
Class 4 15,317 particles 2.3%
Class 5 73,582 particles 11%
Selected classes
Representative 2D classes
Non-uniform refinement (C1)
Reference-based motion correction & CTF correction 512 pix box size
3D Variability Analysis
481,408 particles
3D classification in RELION 10 classes
Representative 3D classes
Class 1 8,041 particles 2%
Class 2 8,149 particles 2%
Class 3 5,588 particles 1.3%
Class 4 32,451 particles 7.4%
Class 5 422,989 particles 86.6%
Focussed 3D classification 10 classes
90°
Class 1 59,251 particles 12.3%
Class 2 58,851 particles 12.2%
Class 3 68,644 particles 14.3%
Class 4 64,474 particles 13.4%
Class 5 30,282 particles 6.3%
Class 6 52,528 particles 11%
Class 7 22,974 particles 4.8%
Class 8 32,442 particles 6.8%
Class 9 36,271 particles 7.5%
Class 10 55,727 particles 11.6%
Non-uniform refinement (C1)
Focused refinement
2.5 4.5 6.5
Local resolution (Å)
Gold-standard FSC curves
Viewing direction distribution
phenix. combine_focused_maps
180°
Class 1
Class 2
Class 3
Class 4

**Supplemental figure 4. Cryo-electron microscopy data processing pipeline.** Representative micrographs and representative 2D classifications of the PafA-ADP-PanB~Pup quaternary complex are shown. 3D refinement and classifications were performed, followed by focused refinements for the PafA and PanB components of the maps. Gold-standard Fourier shell correlation (FSC) curves and plots are displayed. The final combined focused maps are shown for quaternary complex classes 1-4.

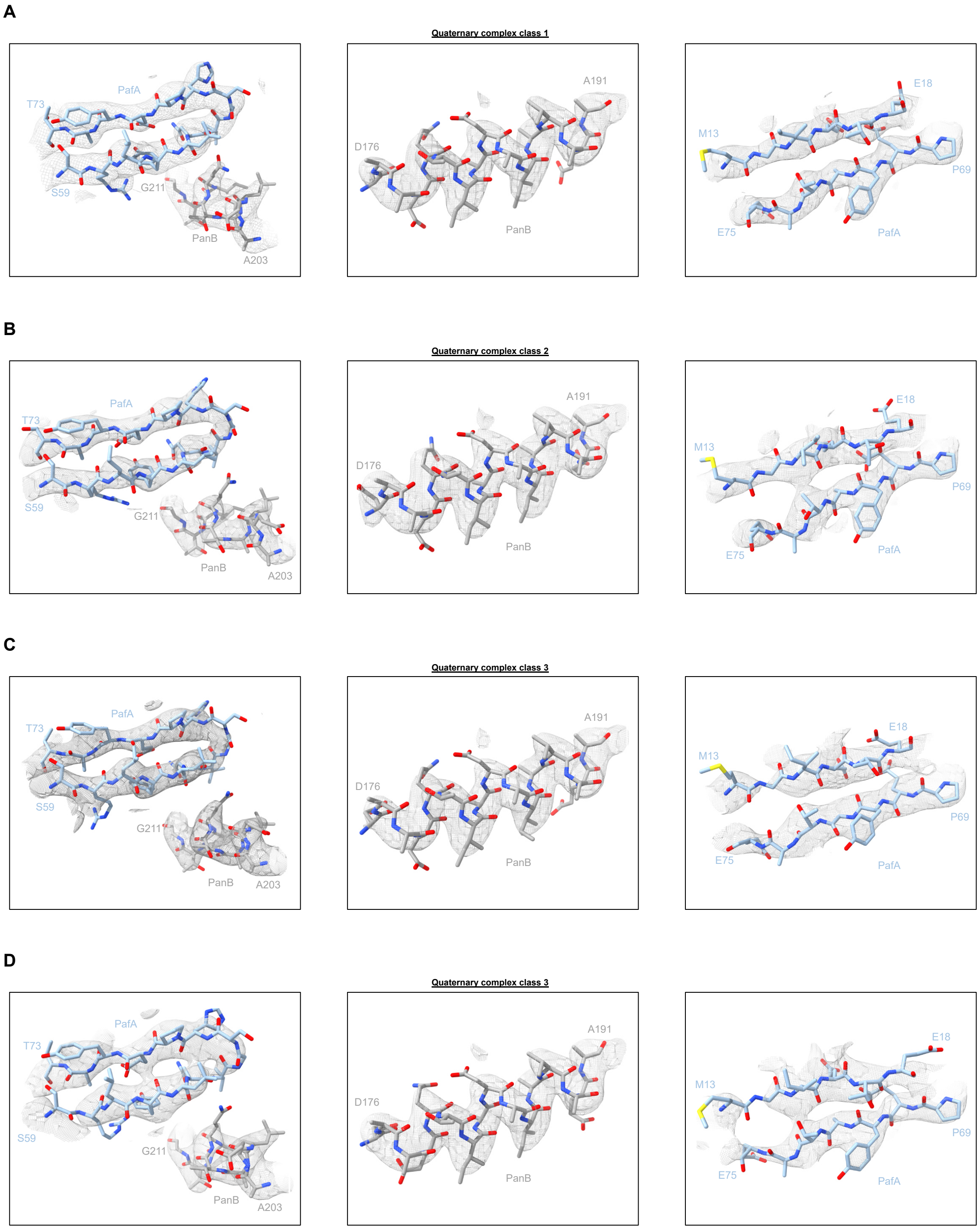


**Supplemental figure 5. Model to map fit for representative regions.** Selected regions of PafA and PanB are shown for **A.** class 1, **B.** class 2, **C.** class 3, and **D.** class 4.

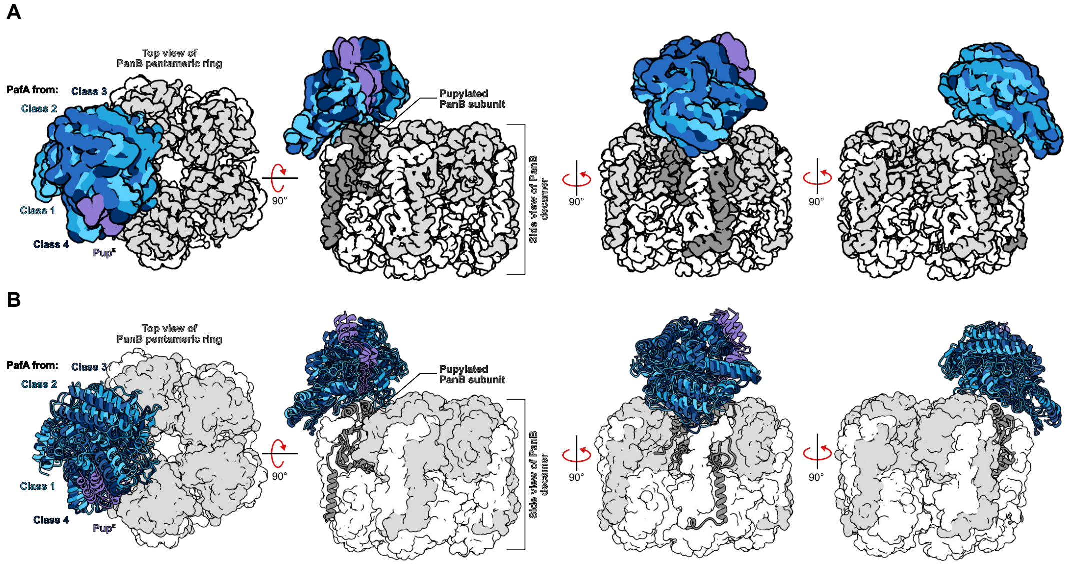


**Supplemental figure 6. Cryo-EM reveals different modes of PanB engagement.** Alignment of the four **A.** cryo-EM map and **B.** model classes of the PafA-ADP-PanB~Pup quaternary complex. Pup (purple) occupies the Pup-binding groove of PafA (class 1-4, light to dark blue), while PafA engages the apical surface of PanB (grey). Comparison of the four classes reveals subtle variations in how PafA contacts PanB, suggesting multiple binding modes.

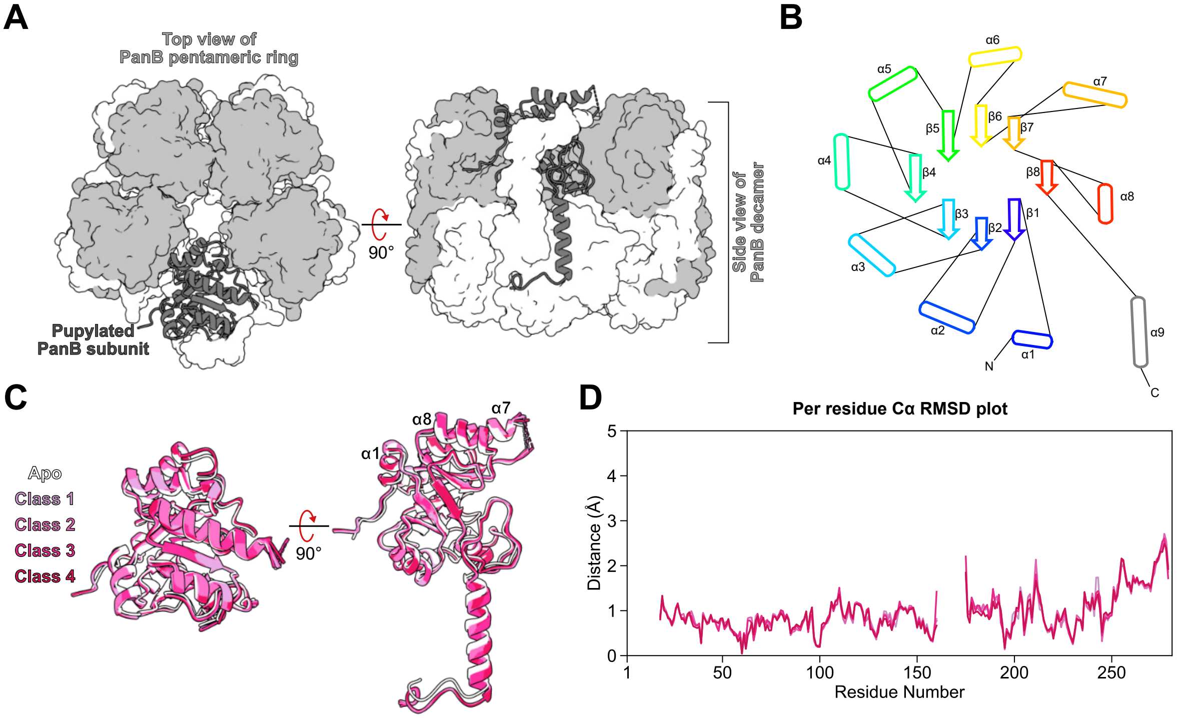


**Supplemental figure 7. PanB adopts similar conformations in apo and PafA-bound states.** **A.** Top and side views of PanB decamer from the PafA-ADP-PanB~Pup quaternary complex from class 1. One pentameric ring is shown as a white surface, while the other is light grey. The pupylated PanB subunit that gets engaged by PafA is shown as a dark grey cartoon. PanB from the apo crystal structure [PDB: 1OY0 (*28*)] and from the four quaternary classes presented here, adopt this canonical decamer formed by the stacking of two pentameric rings. **B.** Two-dimensional topology map of a PanB subunit showing a modified $(\beta/\alpha)_8$-barrel extended by a C-terminal α9 helix, generated with 2DProts (*57*) and modified. **C.** Alignment of one subunit from the apo PanB crystal structure [PDB: 1OY0 (*28*), white] and the pupylated PanB subunit from the four quaternary complex classes (light to dark pink). The Cα RMSD across all atom pairs for PanB from the crystal structure [PDB: 1OY0 (*28*)] to the pupylated PanB subunit from the four quaternary complexes is 1.03, 1.04, 1.04, and 1.02 Å, respectively, indicating minor change in the overall architecture of PanB. **D.** Per residue Cα RMSD plot from the alignment in C. Sequence spanning unmodeled regions are shown as gaps.

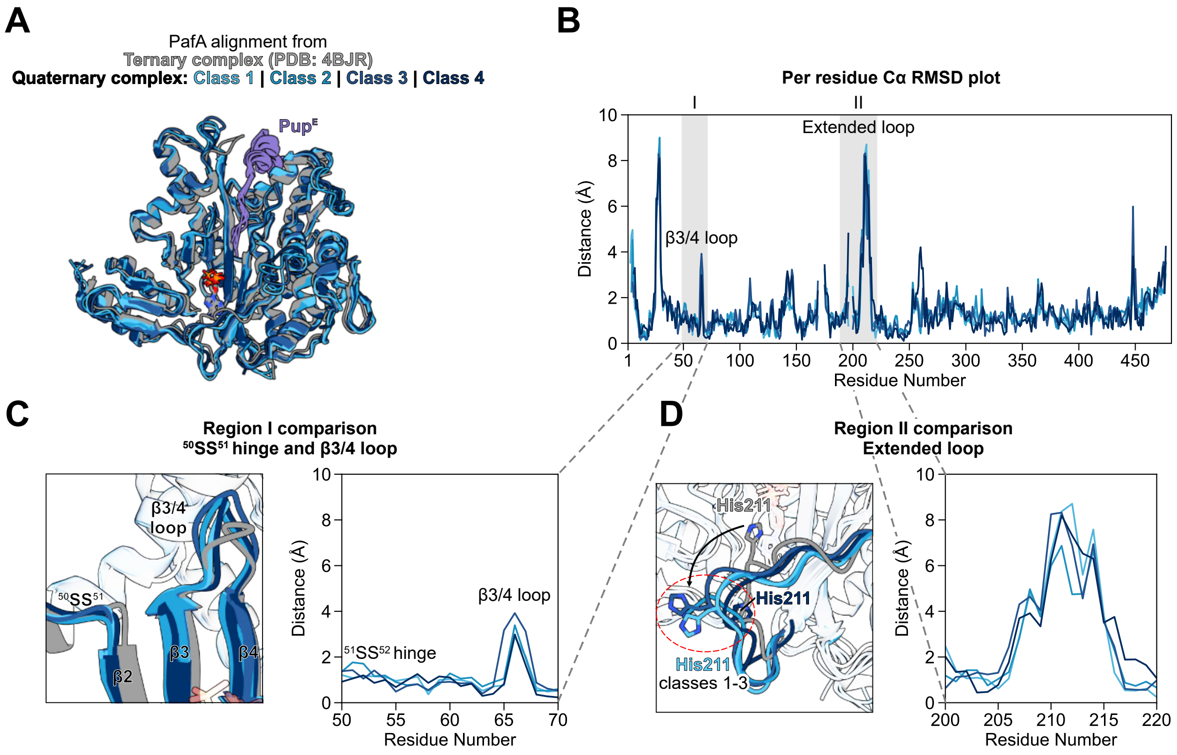


**Supplemental figure 8. Regions involved in PanB engagement undergo conformational changes in PafA. A.** Alignment of PafA from the four quaternary complex classes (light to dark blue) against PafA from the ternary complex crystal structure [PDB: 4BJR (*16*) , grey]. Pup is shown in purple, occupying the Pup-binding groove, and nucleotide (ATP – crystal structure, ADP – cryoEM classes) in grey. The Cα RMSD across all atom pairs for PafA from the crystal structure [PDB: 4BJR (*16*) ] to PafA from the four quaternary complexes is 1.68, 1.69, 1.77, and 1.74 Å, respectively, indicating minor change in the overall architecture of PafA. **B.** Per residue Cα RMSD plot from the alignment in A. Regions I ($^{50}$SS$^{51}$ hinge and β 3/4 loop) and II (extended loop) involved in PanB engagement are highlighted in grey. Sequence spanning unmodeled regions are shown as gaps. **C.** and **D.** Insets for regions I ($^{50}$SS$^{51}$ hinge and β 3/4 loop) and II (extended loop), respectively. PanB engagement induces changes in residues within the β3/4 loop and extended loop of PafA, highlighted key differences across the four quaternary complex classes.

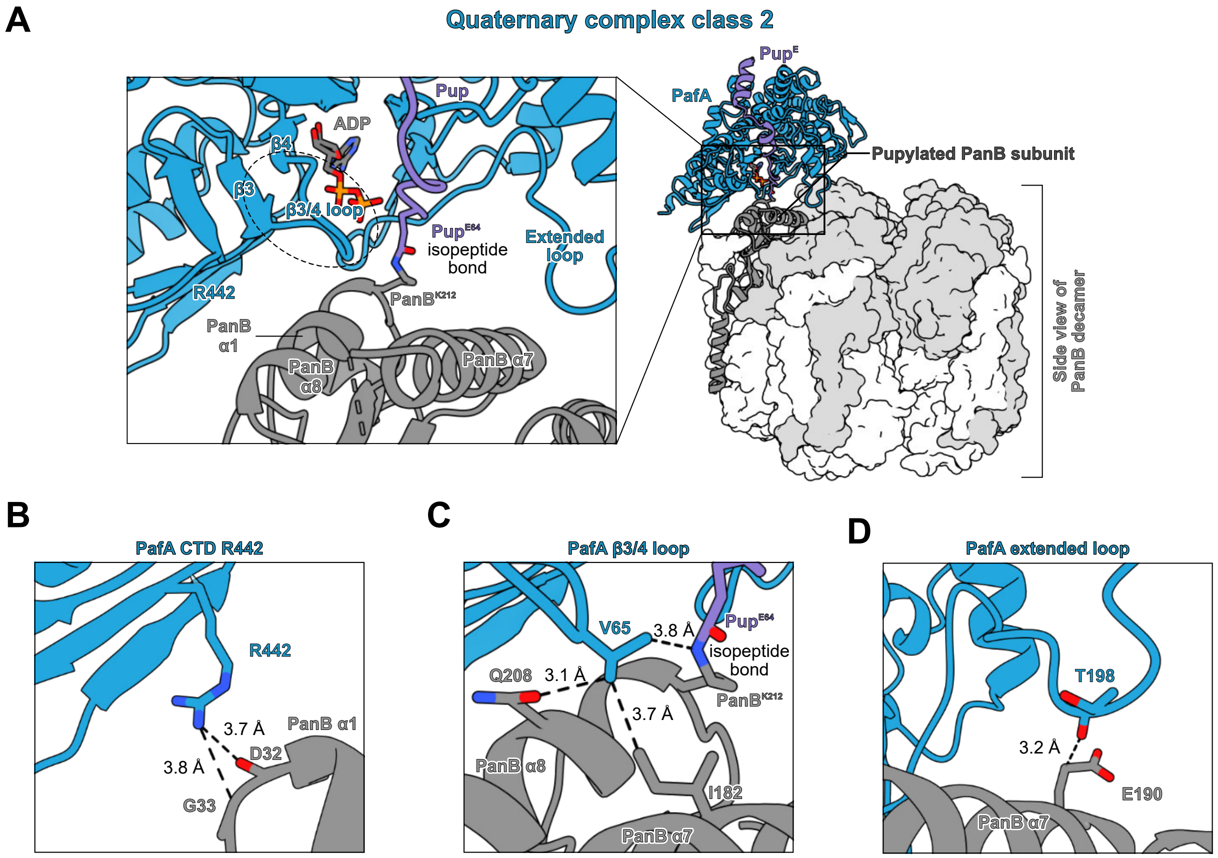


**Supplemental figure 9. PafA-PanB contacts in quaternary complex class 2. A.** The refined model of the class 2 quaternary complex is shown. PafA (blue), Pup (purple), and the pupylated PanB subunit (dark grey) are depicted as cartoons, while the remaining nine PanB subunits are rendered as a surface. PafA engages Pup within the Pup-binding groove and simultaneously contacts the apical surface of the pupylated PanB subunit. The inset highlights the PafA-PanB interface, with the three apical helices of PanB (α1, α7, and α8, the latter containing the pupylated K212) indicated. The isopeptide bond between PanB K212 and Pup E64 is shown, and PafA's β3-4 loop, extended loop, and CTD residue R442, involved in target protein engagement are highlighted. For clarity, PafA α1 (residues 33-46) is omitted from the inset. **B.** Interaction between PafA R442 and PanB D32 3.7 Å (NH2-O), and G33 3.8 Å (NH2-Cα). **C.** Interactions between PafA V65 and PanB I182 3.7 Å (Cγ 1-Cδ1), Q208 3.1 Å (Cγ 1-Oε1), and pupylated K212 3.8 Å (Cγ 2-NZ). **D.** Interaction between PafA T198 and PanB E190 3.2 Å (O-Cγ).

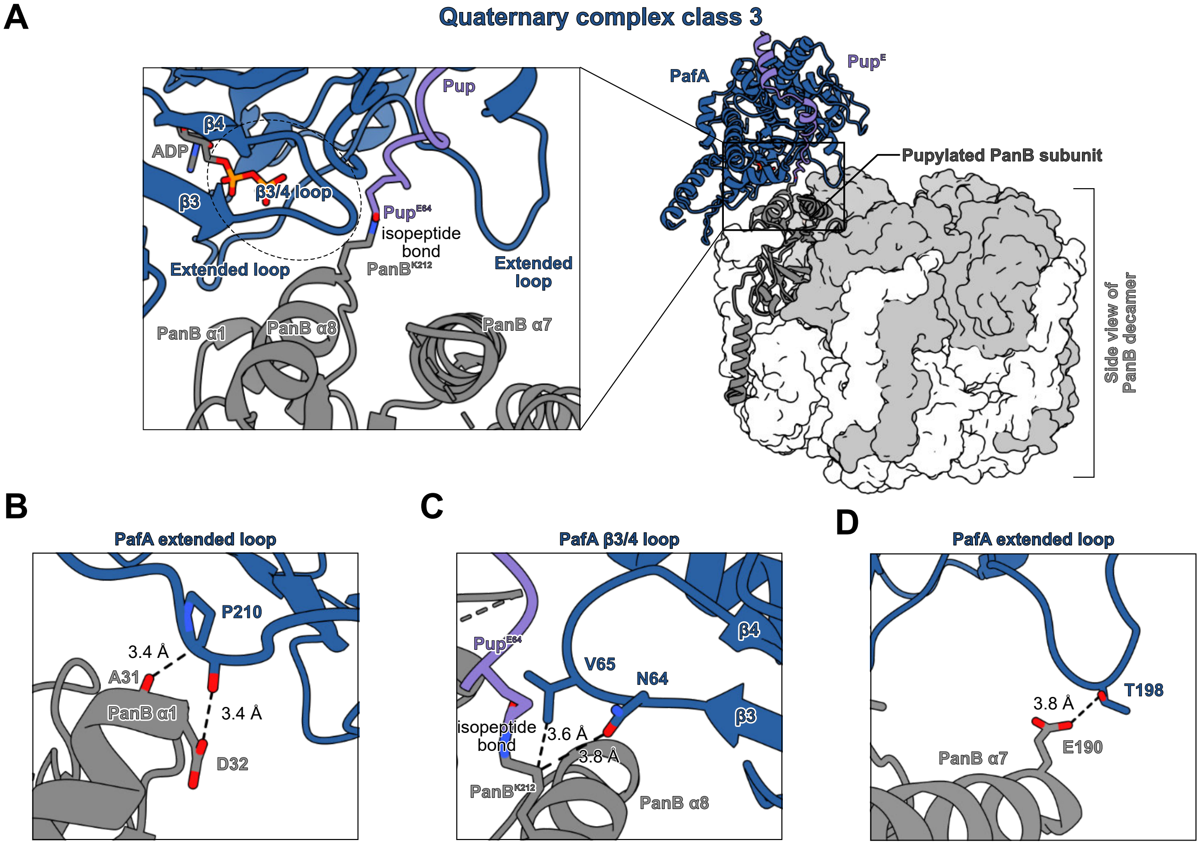


**Supplemental figure 10. PafA-PanB contacts in quaternary complex class 3. A.** The refined model of the class 3 quaternary complex is shown. PafA (blue), Pup (purple), and the pupylated PanB subunit (dark grey) are depicted as cartoons, while the remaining nine PanB subunits are rendered as a surface. PafA engages Pup within the Pup-binding groove and simultaneously contacts the apical surface of the pupylated PanB subunit. The inset highlights the PafA-PanB interface, with the three apical helices of PanB (α1, α7, and α8, the latter containing the pupylated K212) indicated. The isopeptide bond between PanB K212 and Pup E64 is shown, and PafA's β3-4 loop and extended loop, involved in target protein engagement are highlighted. For clarity, PafA α1 (residues 33-46) is omitted from the inset. **B.** Interactions between PafA P210 and PanB A31 3.4 Å (Cβ-O), and D32 3.4 Å (O-Oδ1). **C.** Interaction between PafA N64 and PanB K212 3.8 Å (Oδ1-Cδ), and PafA V65 with PanB K212 3.6 Å (Cγ2-Cδ). **D.** Interaction between PafA T198 and PanB E190 3.8 Å (Oγ 1-Oε1).

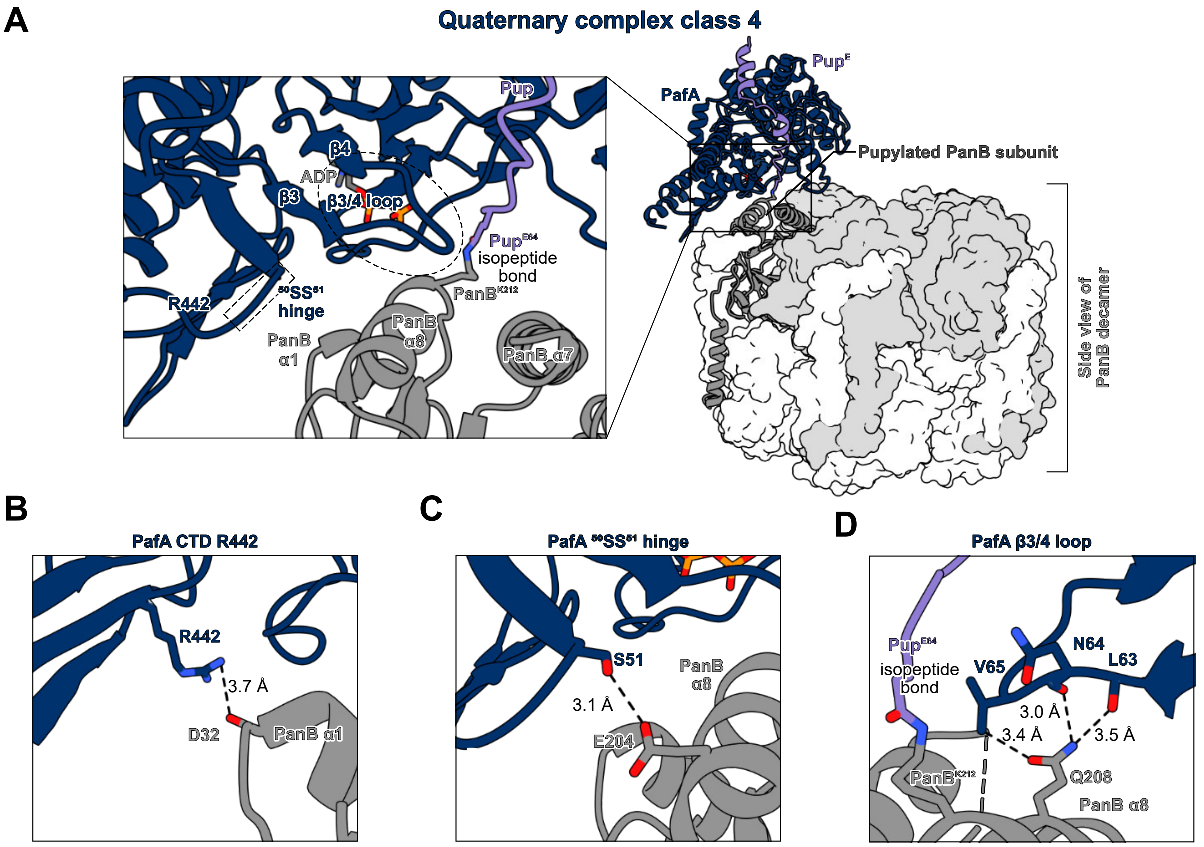


**Supplemental figure 11. PafA-PanB contacts in quaternary complex class 4. A.** The refined model of the class 4 quaternary complex is shown. PafA (blue), Pup (purple), and the pupylated PanB subunit (dark grey) are depicted as cartoons, while the remaining nine PanB subunits are rendered as a surface. PafA engages Pup within the Pup-binding groove and simultaneously contacts the apical surface of the pupylated PanB subunit. The inset highlights the PafA-PanB interface, with the three apical helices of PanB (α1, α7, and α8, the latter containing the pupylated K212) indicated. The isopeptide bond between PanB K212 and Pup E64 is shown, and PafA's $^{50}$SS$^{51}$ hinge, β3-4 loop, and CTD residue R442, involved in target protein engagement are highlighted. For clarity, PafA α1 (residues 33-46) is omitted from the inset. **B.** Interaction between PafA R442 and PanB D32 3.7 Å (NH1-O). **C.** Interaction between PafA S51 and PanB E204 3.1 Å (Oγ -Oε1). **D.** Interactions between PafA L63 and PanB Q208 3.5 Å (O-Nε2), PafA N64 and PanB Q208 3.0 Å (O-Nε2), and PafA V65 and PanB Q208 3.4 Å (Cγ 1-Oε 1).

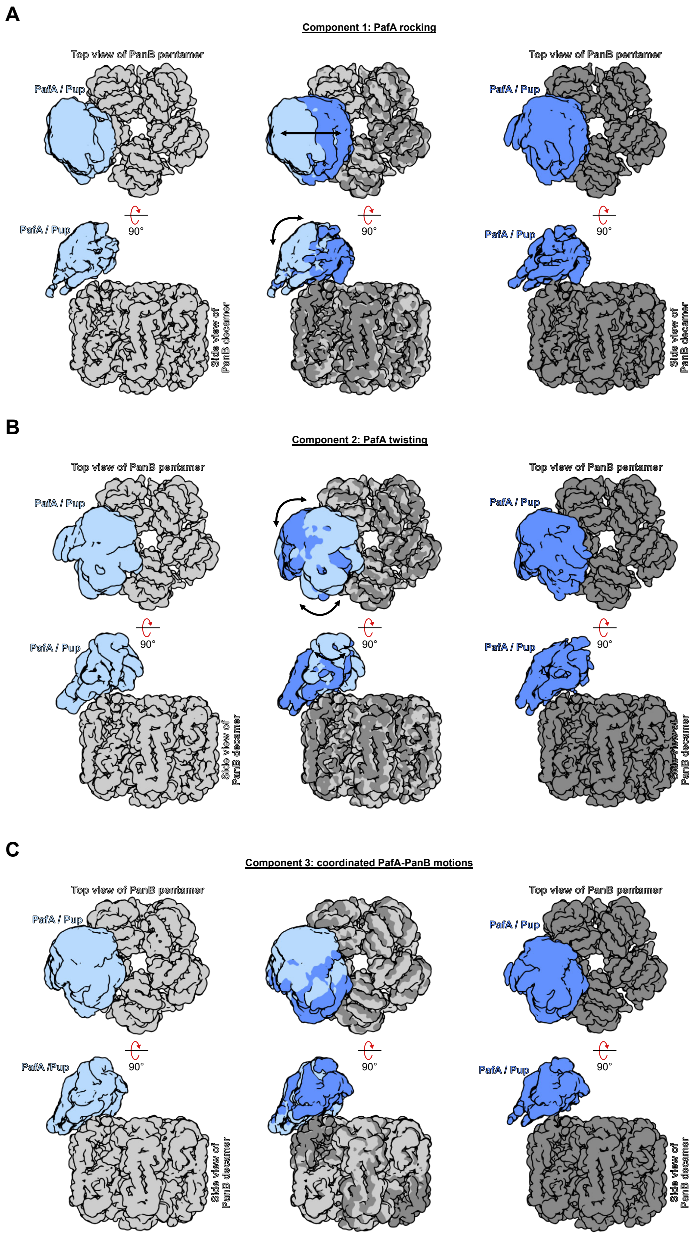


**Supplemental figure 12. 3D variability analysis of the PafA-ADP-PanB~Pup quaternary complex reveals three variable components.** Maps generated at negative (initial frame, PafA/Pup in light blue and PanB in light grey) and positive (final frame, PafA/Pup in dark blue and PanB in dark grey) latent coordinates along the variability components are overlaid for each component. The motions are also shown in Supplementary Movies 2-4. **A.** Captures a rocking motion in PafA, whereas **B.** shows PafA twisting, and **C.** reveals a coordinated movement between PafA and PanB.

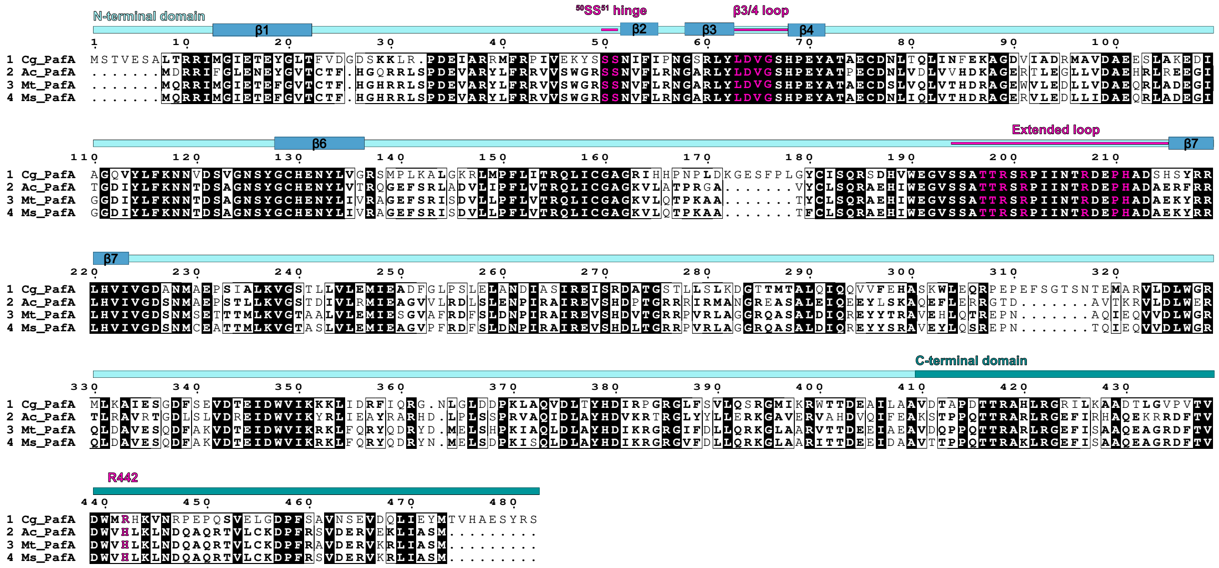


**Supplemental figure 13. Regions involved in target protein engagement are conserved.** A multiple sequence alignment was performed on a curated list of 62 PafA sequences. Residues that are similar between species are bolded, and residues that are strictly conserved are highlighted black. The three regions of PafA (region I - $^{50}$SS$^{51}$ hinge and the β3-4 loop (residues 63-66), region II - extended loop (residues 196-216), and region III - R442 from the C-terminal domain) that are involved in target protein recruitment are in pink text. Residues in the hinge, β3-4 loop, and extended loop are strictly conserved across all PafA sequences. While R442 is not strictly conserved, the positive charge is retained across 59/62 PafA sequences (R: 15%, H: 80%, T: 5%).

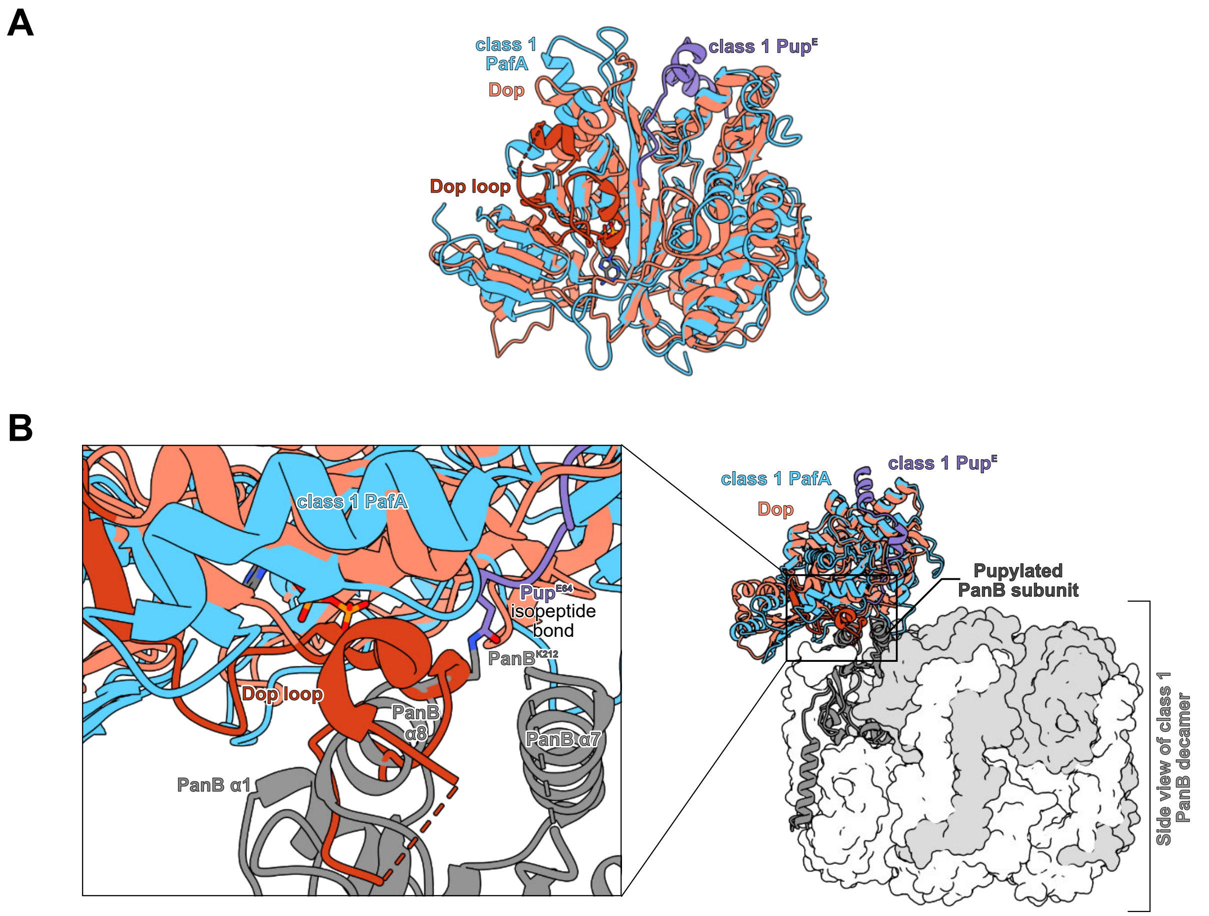


**Supplemental Figure 14. Alignment of Dop supports a negative regulatory role for the Dop loop. A.** Alignment of structural homologs PafA (quaternary complex 1, light blue) and Dop [PDB: 7OXV (*13*), salmon]. The Dop loop (dark salmon, residues 36-80) is unique to Dop and is positioned towards the conserved active site β-sheet cradle. **B.** Alignment from A in the context of the full PafA-ADP-PanB~Pup quaternary complex reveals how Dop may engage pupylated PanB. The inset shows the Dop loop occupies the same space as PanB's pupylated α8 helix, suggesting possible interference and supporting its role as a negative regulator of depupylation.
